# Mitigation of Stress-induced Structural Remodeling and Functional Deficiency in iPSC-CMs with PLN R9C Mutation by Promoting Autophagy

**DOI:** 10.1101/2024.04.17.589921

**Authors:** Qi Yu, Robert J Barndt, Yawei Shen, Karim Sallam, Ying Tang, Stephen Y. Chan, Joseph C. Wu, Qing Liu, Haodi Wu

**Affiliations:** Pittsburgh Heart, Lung, and Blood Vascular Medicine Institute, University of Pittsburgh School of Medicine, Pittsburgh, PA 15261, USA; Department of Biological Sciences, Clemson University, Clemson, SC 29634, USA; Center for Human Genetics, Clemson University, Greenwood, SC 29646, USA; Stanford Cardiovascular Institute, Department of Medicine, Division of Cardiovascular Medicine, and Institute for Stem Cell Biology and Regenerative Medicine, Stanford University School of Medicine, Stanford, CA 94304, USA; Division of Cardiology, Department of Medicine, University of Pittsburgh School of Medicine, Pittsburgh, PA 15261, USA

## Abstract

**Background:** Phospholamban (PLN) is a key regulator of cardiac function connecting adrenergic signaling and calcium homeostasis. The R9C mutation of PLN is known to cause early onset dilated cardiomyopathy (DCM) and premature death, yet the detailed mechanisms underlie the pathologic remodeling process are not well defined in human cardiomyocytes. The aim of this study is to unravel the role of PLN R9C in DCM and identify potential therapeutic targets.

**Methods:** PLN R9C knock-in (KI) and patient-specific induced pluripotent stem cell-derived cardiomyocytes (iPSC-CMs) were generated and comprehensively examined for their expression profile, contractile function, and cellular signaling under both baseline conditions and following functional challenges.

**Results:** PLN R9C KI iPSC-CMs exhibited near-normal morphology and calcium handling, slightly increased contractility, and an attenuated response to β-adrenergic activation compared to wild-type (WT) cells. However, treatment with a maturation medium (MM) has induced fundamentally different remodeling in the two groups: while it improved the structural integrity and functional performance of WT cells, the same treatment result in sarcomere disarrangement, calcium handling deficiency, and further disrupted adrenergic signaling in PLN R9C KI cells. To understand the mechanism, transcriptomic analysis showed the enrichment of protein homeostasis signaling pathways specifically in PLN R9C KI cells in response to the MM treatment and increased contractile demands. Further studies also indicated elevated ROS levels, interrupted autophagic flux, and increased pentamer PLN aggregation in functionally challenged KI cells. These results were further confirmed in patient-specific iPSC-CM models, suggesting that functional stresses exacerbate the deficiencies in PLN R9C cells through disrupting protein homeostasis. Indeed, treating stressed patient cells with autophagy-accelerating reagents, such as metformin and rapamycin, has restored autophagic flux, mitigated sarcomere disarrangement, and partially rescued β-adrenergic signaling and cardiac function.

**Conclusions:** PLN R9C leads to a mild increase of calcium recycling and contractility. Functional challenges further enhanced contractile and proteostasis stress, leading to autophagic overload, structural remodeling, and functional deficiencies in PLN R9C cardiomyocytes. Activation of autophagy signaling partially rescues these effects, revealing a potential therapeutic target for DCM patients with the PLN R9C mutation.

**Graphic abstracts:** A graphic abstract is available for this article.

## Introduction

During the contraction of cardiac muscle (systole), the depolarization of the membrane triggers calcium influx through activation of L-type calcium channels (LCC). The increase of cytosol calcium concentration induces the opening of ryanodine receptors (RYR2) and the release of a large amount of calcium from intracellular stores in the sarcoplasmic reticulum (SR), a mechanism known as calcium induced calcium release (CICR).^1^ The released calcium will bind to the troponin complex in the sarcomere contractile apparatus and initiate the contraction of cardiac cells. During the relaxation of cardiac muscle (diastole), calcium is removed from the cytosol and recycled into the SR by the sarco/endoplasmic reticulum Ca^2+^-ATPase (SERCA), a calcium pump embedded in the SR membrane. The calcium affinity of SERCA is negatively regulated by a small single-pass membrane protein phospholamban (PLN).^2^ Upon β-adrenergic signaling activation in the heart, PLN can be phosphorylated by cAMP dependent protein kinases A (PKA) and calcium/calmodulin-dependent protein kinase II (CaMKII) at Ser 16 and Thr 17, respectively. The phosphorylation promotes the conformational switch of PLN, which reverses the binding and inhibition of PLN on SERCA, allowing SERCA to pump calcium ions more rapidly into the SR.^3^ This results in increased SR calcium recycling (lusitropy), faster beating rate (chronotropy), and enhanced contractility (inotropy), thus releasing the reserved pumping power of the heart in response to higher functional demand. Therefore, PLN is a key regulatory molecule that connects calcium handling and β-adrenergic signaling pathways in the heart.^4^

Given the important regulatory role of PLN in the heart, PLN mutations significantly affect normal cardiac function. Indeed, previous studies have identified several PLN mutations in families of patients with dilated cardiomyopathy (DCM), a more common disorder that leads to heart failure (HF). These PLN mutations include R14del, R9C, R25C, L39X, E2X and others.^5–11^ Among these mutations, R14del, R9C and R25C are the most common in human DCM. Studies have shown that PLN-R14del mutation results in impaired contractile function, increased myocardial fibrosis, PLN protein aggregation, and increased risk of ventricular tachycardia (VT).^12–14^ Also, PLN-R25C mutation is known to cause the super-inhibition of SERCA2a, decreased SR calcium load, increased SR calcium leak and CaMKII activity, which result in weaker contraction and higher risk of arrhythmia.^7, 15^ PLN-R9C was first identified by Schmitt *et al.* in patients with early onset DCM, who hypothesized the mutated PLN-R9C protein binds irreversibly to the catalytic subunit of PKA (PKA-C), which prevents phosphorylation of PLN at Ser 16.^9^ Moreover, a PLN R9C animal model was created to understand the pathological mechanism of PLN R9C, which confirmed defective β-adrenergic signaling in the heart.^16, 17^ Other studies also indicate the PLN R9C mutation affects the binding affinity of PLN to SERCA and the monomer-pentamer equilibrium of PLN protein.^18, 19^ However, due to the lack of human patient samples, the role of PLN R9C mutation in human DCM remains elusive, and there is no specific treatment for DCM patients with this mutation to date.

To better understand the pathological role of PLN R9C mutation in human cardiomyocytes, the current study generated induced pluripotent stem cells (iPSCs) disease models with CRISPR/Cas9 edited isogenic knock-in (KI) and DCM patient-specific lines that carry PLN R9C mutation. The iPSC-derived cardiomyocytes (iPSC-CMs) carrying PLN R9C mutation exhibited near normal calcium handling and contractility features at baseline, but recapitulated the blunted β-adrenergic signaling as reported in animal models and human cell studies.^20^ After treatment of maturation medium (MM), which promotes metabolic and functional maturity, WT cells showed enhanced maturation and functional performance while functional deficiency was observed in PLN R9C groups, indicating pathological remodeling upon increased contractile demand in mutation-carrying cells. Further transcriptome profiling and signaling studies indicated ER/autophagic stress and protein aggregation were exacerbated in PLN R9C cells after MM treatment, which were accompanied by an increased level of reactive oxygen species (ROS) and sarcomere disarrangement. Finally, our study showed activation of autophagic signaling with metformin and rapamycin can partially restore morphological and functional integrity in patient-specific iPSC-CMs.

In summary, the current study has demonstrated the critical role of protein homeostasis signaling during the pathogenesis of PLN R9C mutation-induced DCM and has identified novel therapeutic targets.

## Methods

### Recruitment of dilated cardiomyopathy patient with PLN R9C mutation

A male patient (age = 44) diagnosed with severe dilated cardiomyopathy and markedly depressed cardiac function (EF 25∼35%) was recruited and consented to the current study. The recruitment of patient tissue samples followed the IRB#29904 of Stanford University, approved by the SU ethics committee. Skin biopsy was isolated and rinsed in PBS. The skin biopsy was then minced and digested in DMEM medium (Dulbecco’s modified Eagle medium) containing 1mg/ml collagenase I at 37°C for 6 hours. Dermal fibroblasts were collected by centrifugation at 300 × g for 5 min, and then maintained with DMEM containing 10% FBS, 100 U/ml Penicillin, and 100 U/ml Streptomycin. The fibroblasts were used for reprogramming within 5 passages.

### Reprogramming of patient-specific iPSCs

Patient-specific iPSCs were generated from fibroblasts using CytoTune-iPS 2.1 Sendai Reprogramming Kit (Thermo Fisher) following the manufacturer’s instructions. Briefly, patient dermal fibroblasts were maintained in DMEM + 10% FBS and reseeded on a 6-well plate at a density of 1∼2×10^5^ cells per well 2 days before virus transduction. When cells are 30∼60% confluent, Sendai Virus mixture (expressing KOS, hL-Myc, hKlf4) was applied to the cell culture with MOIs of 5 for KOS, 5 for hL-Myc, and 3 for hKlf4. The culture medium was be replaced with xeno-free fibroblast medium 24 h after transduction. The cells will was in the same well before passaging. After seven days of transduction, fibroblast cells were harvested and seeded to Matrigel-coated 6-well plates. After 24 h, the culture media was changed to Essential 8 (E8) (Giboco), which was replaced every day until single iPSC clones were observed in the plate. Picked clones were kept in culture and passaged at least 10 generations. The quality of these iPSCs was validated by immuno-staining of pluripotent markers such as SSEA-4, Nanog, SOX2, and OCT4 as well as in vitro 3-germ layer differentiation assays with STEMdiff™ Trilineage Differentiation Kit (StemCell). The successful differentiation of ectoderm, endoderm, and mesoderm was confirmed by immunostaining of γ-tubulin, SOX17, and brachyury.

### Plots and statistical Analysis

All data in this study are presented as mean ± SEM from at least three independent experiments. Statistical analyses were calculated using unpaired two-tailed Student’s *t* test to compare two normally distributed data sets. One-way or two-way ANOVA of variance tests followed by Holm-Sidak method were used for multi-group comparisons of data. *p* < 0.05 was regarded as significant. The statistical analyses were performed using GraphPad Prism 9.

### Data availability

Detailed experimental methods are available in the Supplemental Material. Key research materials are listed in the major resources table in the Supplemental Material. The data presented in this study is available on request from the corresponding author. The raw data from RNA-seq was uploaded to the NIH BioProject database at https://www.ncbi.nlm.nih.gov/bioproject. (ID: PRJNA1045604)

## Results

### PLN R9C KI iPSC-CMs show blunted adrenergic signaling

A heterozygous PLN R9C mutation was introduced into wild type (WT) iPSC lines using CRISPR tools as previously described.^21^ WT and PLN R9C KI (GE) iPSC lines were differentiated into beating cardiomyocytes with standard protocols.^22^ Both flow cytometry analysis (FACS) of cardiac troponin T (TNNT2)-positive cells and immunostaining of cardiac markers, including TNNT2 and alpha-actinin, consistently indicated that the PLN R9C mutation did not adversely affect the efficiency and purity of cardiomyocyte differentiation. (**Supplementary Figure 1A-1D**) All the structural and functional analyses of WT and PLN R9C-carrying iPSC-CMs in the current study were performed in purified iPSC-CM culture medium for 35∼50 days after differentiation.

We first examined the functional performance of both WT and PLN R9C KI iPSC-CMs with Fluo-4 AM calcium handling imaging (**Figure 1A**). To compare β-adrenergic signaling in both WT and GE cells, 100 nM isoproterenol (ISO) was applied to the iPSC-CMs (**Figure 1A**). Our result showed WT and PLN R9C KI iPSC-CMs have similar calcium handling properties such as beating rate and calcium transient time to peak at basal level (**Figure 1B** and **1C**), except for a slightly increased calcium recycling rate in terms of shorter decay Tau (249.7 ± 9.2 ms vs. 206.5 ± 8.3 ms) in the R9C KI cells (**Figure 1D**). We also measured the contractility of the iPSC-CMs using traction force microscopy (**Figure 1E**), which showed the relaxation velocity of mutant cells was slightly increased in mutant cells compared to the WT group at basal level (5.68 ± 0.30 um/s vs. 7.62 ± 0.19 um/s), while no significant changes in the contractile velocity was observed (**Figure 1F** and **1G**). The treatment of ISO significantly accelerated calcium recycling (249.7 ± 9.2 ms vs. 166.6 ± 9.3 ms) (**Figure 1D**), increased beating rate (29.1 ± 0.9 bpm vs. 74.4 ± 2.4 bpm) (**Figure 1C**), and enhanced contraction (9.55 ± 0.31 um/s vs. 13.91 ± 0.84 um/s) and relaxation (5.68 ± 0.30 um/s vs. 9.05 ± 0.36 um/s) in the WT iPSC-CMs (**Figure 1F**). However, the same treatment only resulted in a much milder functional response in the mutant iPSC-CMs, such as the faster beating rate (from 29.7 ± 0.6 bpm to 50.3 ± 0.5 bpm, increased by ∼69%) and calcium elevation phase (from 467.9 ± 11.8 ms to 391.2 ± 12.6 ms, decreased by ∼17%), while the calcium transient decay tau and contractility were almost unchanged (**Figure 1F** and **1G**). As a result, the functional performance of WT and GE iPSC-CMs became significantly different after ISO treatment. To understand the contribution of PLN R9C mutation to the β-adrenergic signaling in both WT and GE cells, we measured the phosphorylation of PLN at Ser16 before and after ISO treatment by western blot (**Supplementary Figure 1E**). Our result showed the total PLN level was lower in mutant cells, although the PLN R9C protein still response to the ISO treatment, the phosphorylation of the PLN protein was significantly prohibited by the R9C mutation, both before and after ISO challenge (**Supplementary Figure 1F** and **1G**). All these observations suggest a blunted β-adrenergic signaling in the PLN R9C cells, which is likely due to the resistance of mutant PLN to the phosphorylation by PKA.^23^

**Figure 1.**
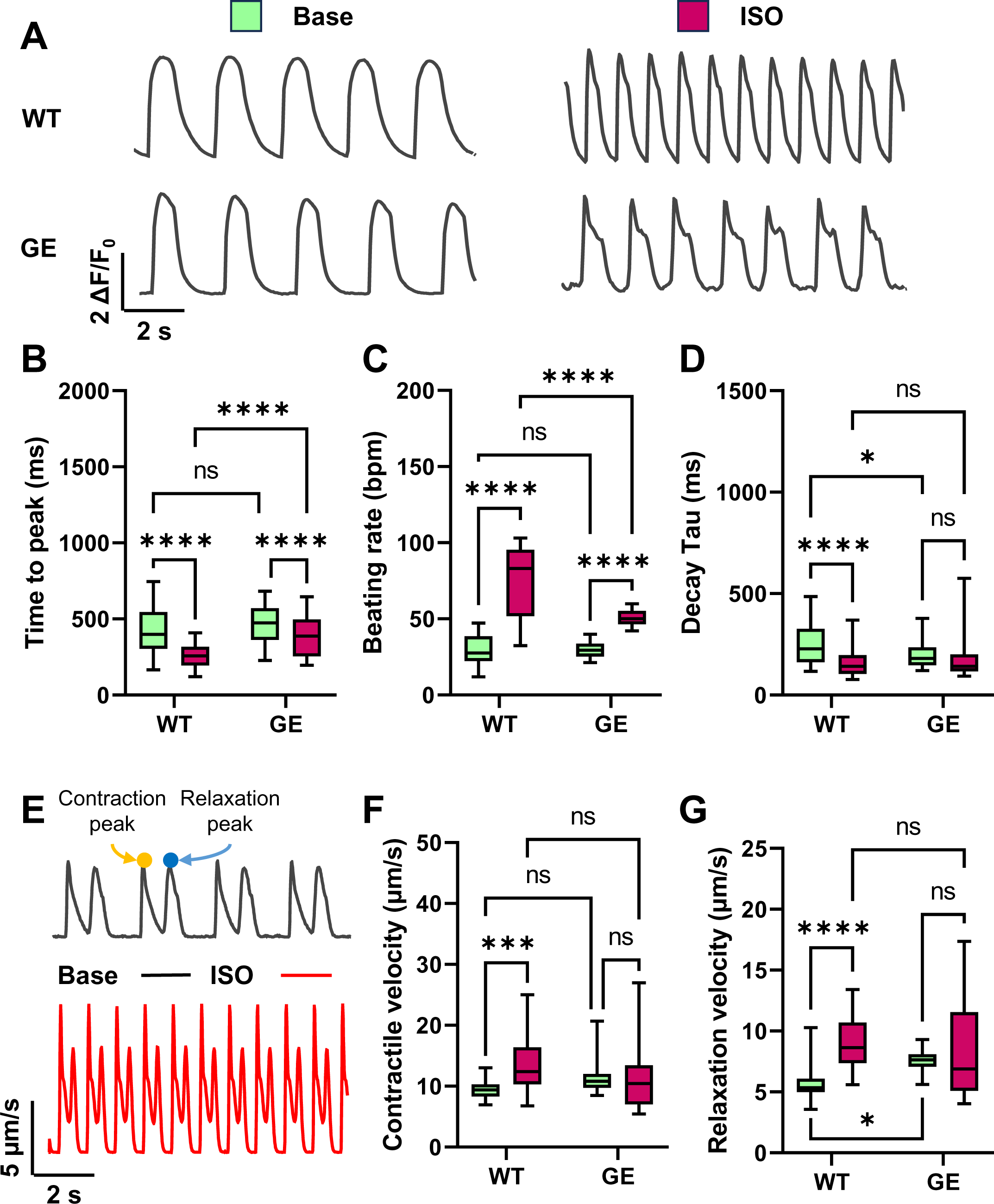
Impaired adrenergic signaling in PLN R9C iPSC-CMs. A. Representative Fluo-4 AM calcium transient trace in WT and PLN R9C (GE) iPSC-CMs before and after isoproterenol (ISO) treatment. B to D. Quantification of calcium handling parameters: transient rise time (B), beating rate (C), and transient decay Tau (D) from WT and GE groups at base level and after ISO treatment. N > 106 cells of WT, GE, WT ISO, and GE ISO groups from at least 3 independent experiments. E. Representative traction force microscopy (TFM) traces of the displacement velocity of beating iPSC-CMs before and after ISO treatment. Yellow and blue arrows indicate the contraction peak and relaxation peak in a beating episode. F and G. Quantification of the peak velocities of contraction (F) and relaxation (G) of WT and GE iPSC-CMs at base level and after ISO treatment. N > 25 ROIs of WT, GE, WT ISO, and GE ISO groups from at least 3 independent experiments. NS, no significance, *p<0.05, ***p<0.001, and ****p<0.0001 by 2-way ANOVA test followed by Holm-Sidak method.

### *In vitro* maturation exaggerated the functional deficiency in PLN R9C KI iPSC-CMs

Previous studies on PLN R9C transgenic mice mainly focused on the acute effect of PLN R9C overexpression, which indicated altered dynamics in PLN-SERCA binding and PLN oligomerization.^18^ Moreover, it was demonstrated that mouse models carrying a PLN R9C transgene and 2, 1, or 0 endogenous PLN alleles all developed dilated cardiomyopathy (DCM) at different ages, suggesting the existence of other cardiac toxic mechanisms besides the regulation of SERCA2a inhibition by PLN.^16^ However, previous studies have never examined the pathologic mechanism in PLN R9C-carrying cardiomyocytes under potential pathological stresses. Thus, we sought to understand how functional stresses regulate the functional performance in both WT and PLN R9C KI iPSC-CMs. To do that, we treated iPSC-CMs from both groups with pro-maturation medium (MM) as previously reported.^24^ In our previous studies, we have demonstrated that MM promotes the structural and functional maturation of iPSC-CMs, while at the same time enhances the functional challenges to the iPSC-CMs in all aspects, including metabolism, protein homeostasis, and autophagic flow.^25^ In this study, both WT and PLN R9C KI iPSC-CMs at day 35∼45 of differentiation were maintained in regular medium or treated with MM for additional 7 days to induce maturation and functional stress. Afterwards, we examined the structural integrity of the cells with sarcomere structure immunostaining (**Figure 2A**). The arrangement of the sarcomere structures in each group was further analyzed using a Fast Fourier Transformation (FFT) based algorithm, and the regularity score of the sarcomere protein arrangement was calculated as “power”, and the average length of sarcomere was calculated as “period” (**Supplementary Figure 2A** and **2B**). Our results show that similar power and period of α-actinin distribution in both WT (0.093 ± 0.011, 2.23 ± 0.12 µm) and PLN R9C KI cells (0.078 ± 0.009, 2.15 ± 0.11 µm) at basal level (**Figure 2B** and **2C**).However, after MM treatment, WT cells showed slightly increased regularity of sarcomere protein distribution (0.093 ± 0.01 vs. 0.146 ± 0.016), which is related to better maturity of iPSC-CMs (**Figure 2C**). On the other hand, PLN R9C cells exhibit disarrangement of sarcomere structure, evidenced by much lower power (0.052 ± 0.008) and prolonged sarcomere period (2.83 ± 0.09 µm) (**Figure 2B** and **2C**), indicating significant structural remodeling in mutant cells in response to functional challenges. We also measured the calcium handling (Fura-2 AM) and contractile function of WT and mutant cells after MM treatment (**Figure 2D**). While most of the functional parameters such as calcium transient amplitude, decay Tau, and contractile/relaxation velocity were significantly improved in WT cells through MM treatment, little or no functional improvement was observed in PLN R9C KI cells (**Figure 2E, 2F, Supplementary Figure 2C** to **2F**). Instead, the decreased calcium transient, elevated diastolic calcium level, impaired calcium recycling, and prolonged relaxation phase in MM treated mutant cells all suggested the maturation challenge had exacerbated the functional deficiency in PLN R9C KI iPSC-CMs.

**Figure 2.**
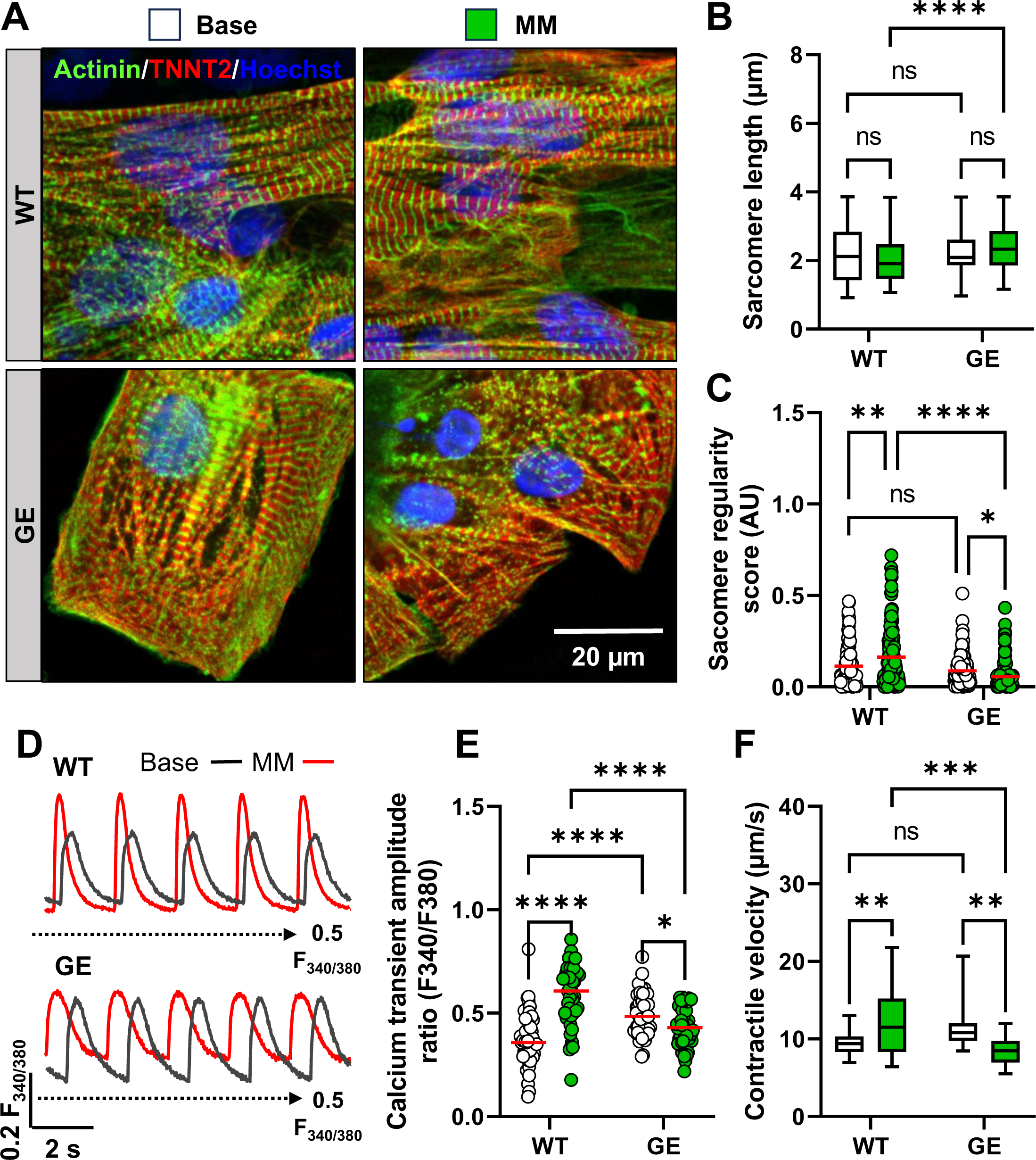
Maturation medium exaggerated the functional deficiency of PLN R9C iPSC-CMs. A. Immunofluorescence staining of WT and GE iPSC-CMs before and after long-term MM treatment with sarcomere markers (green for α-actinin and red for TNTN2). B and C. Fast Fourier transformation (FFT) analysis showed the period (B) and the regularity (C) of α-actinin signal distribution in the iPSC-CMs. MM treatment has improved the regularity of sarcomere arrangement in WT cells, but not in GE cells. N > 78 ROIs of WT, GE, WT MM, and GE MM groups from at least 3 independent experiments. D. Representative ratiometric Fura-2 AM calcium transient trace in WT and GE iPSC-CMs before (black line) and after MM treatment (red line). Black dashed line indicates the F340/F380 ratio of 0.5. E. Long-term MM treatment enhanced Calcium transient amplitude in WT cells, but not in the GE cells. N > 78 cells of WT, GE, WT MM, and GE MM groups from at least 3 independent experiments. F. TFM measurement of contractile velocity in WT and GE cells after MM treatment. N > 25 ROIs of WT, GE, WT MM, and GE MM groups from at least 3 independent experiments. NS, no significance, *p<0.05, **p<0.01, ***p<0.001, and ****p<0.0001 by 2-way ANOVA test followed by Holm-Sidak method.

### RNA-seq revealed that MM treatment activates distinct signaling pathways in WT and PLN R9C KI iPSC-CMs

To better understand the underlying mechanism of the maturation challenge-induced functional deficiency in PLN R9C iPSC-CMs, we performed genome-wide transcriptomic analysis of the differentiated iPSC-CMs from both the WT and mutant group at both basal level and after prolonged MM treatment (**Figure 3A, Supplementary Figure 3A, Supplementary Table 3**). Principal component analysis and inter-sample correlation revealed similar expression patterns and clustering in replicate samples **(Supplementary Figures 3A** and **3B**). We found 618 differentially expressed genes (DEGs) in WT iPSC-CMs before and after MM treatment, while MM led to significant change of expression of 1159 genes in PLN R9C KI cells. Notably, there are only 385 DEGs shared between the 2 groups, indicating the activation of different regulatory mechanisms (**Figure 3A**). The expression profiles of DEGs were extracted and displayed by clustering heatmaps, showing significant differences in the comparison groups **(Figure 3B)**. Indeed, interactive KEGG term analysis revealed that MM treatment induced signaling pathways that are associated with “cardiac muscle contraction”, “protein digestion and absorption”, “oxidative phosphorylation”, and “biosynthesis of amino acids” in WT cells, while the top enriched KEGG terms in PLN R9C KI cells are “protein processing in endoplasmic reticulum”, “protein digestion and absorption”, and “metabolic pathways” (**Figure 3C** and **3D**). Further analysis of the transcriptome showed that the MM induced upregulation of contractile function, calcium handling, and mitochondria relevant genes in WT cells, such as CASQ2, MYL2, MYL3, and COX8A., which is in line with the pro-maturation effects (**Figure 3E**). In contrast, in the MM treated mutant cells, the upregulated genes include: HSPA5, HSPA6, BCL2, HSPA1A and CALR, which are known to regulate protein homeostasis and apoptosis in stressed cardiomyocytes (**Figure 3F**). Moreover, enrichment map profiling of GO biological process terms of the DEGs (Qvalue<0.05) in the WT and mutant groups after MM treatment have revealed the key signaling networks involved in the structural and functional remodeling of the iPSC-CMs in each group, which indicates that functional challenges by MM promoted the cardiac maturity in WT cells but have interrupted protein homeostasis in the PLN R9C KI iPSC-CMs (**Supplementary Figure 3C** and **3D**).

**Figure 3.**
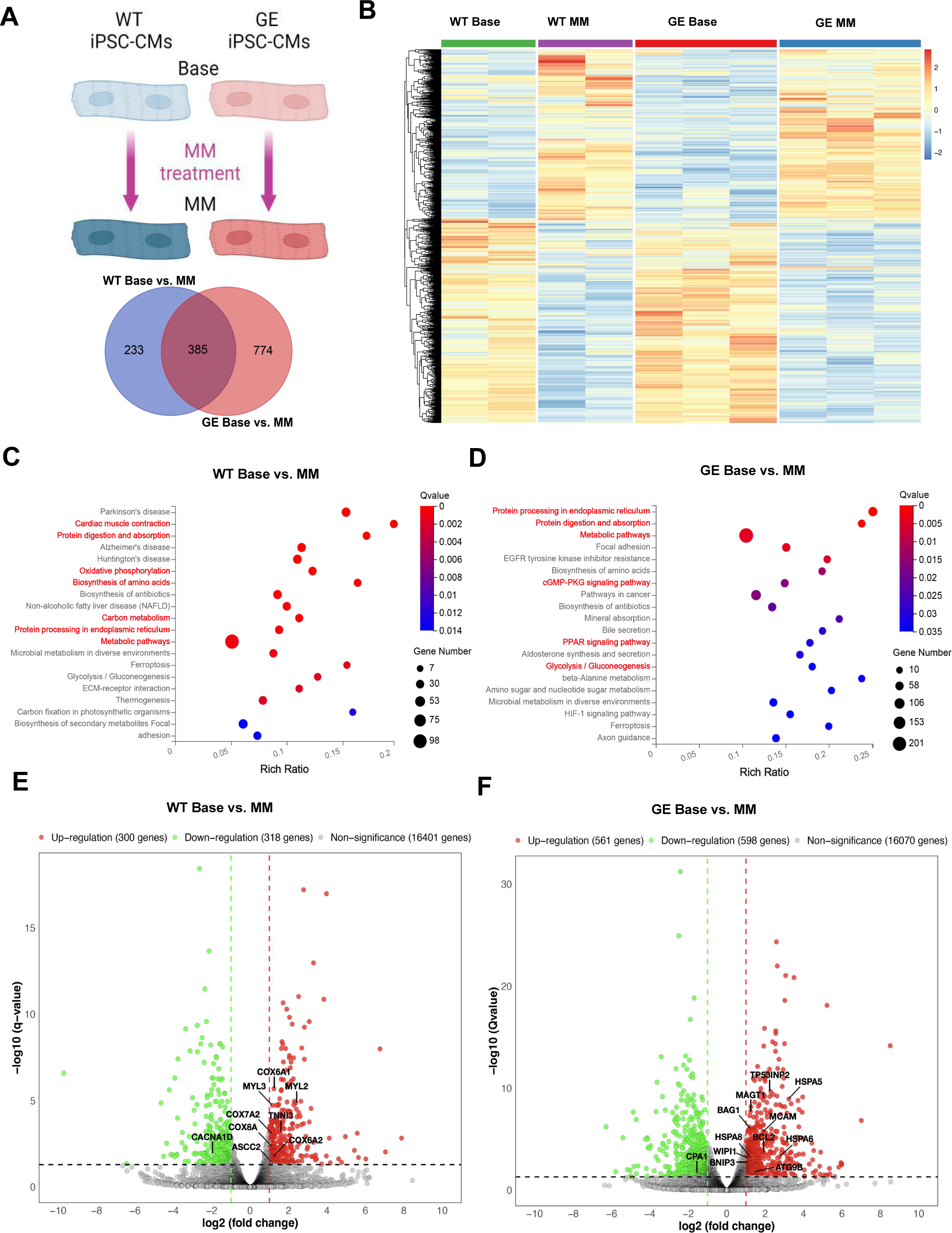
RNA-seq revealed functional challenge interrupt protein homeostasis in PLN R9C iPSC-CMs. A. Comparison of the number of shared and distinct DEGs in WT Base vs. MM and GE Base vs MM comparisons. B. Heatmap indicates the overall DEG expression pattern of all 4 groups of samples: WT Base, WT MM, GE Base, and GE MM. C and D. KEGG analysis highlighted significantly enriched signaling pathways in WT and GE cells in response to MM long-term treatment. E and F. Volcano plot showed the top altered genes after MM treatment in both WT and GE cells. Some representative genes are highlighted in black in the plot.

Next, we validated the expression of DEGs highlighted by RNA-seq data by qPCR (**Figure** 4, **Supplementary Figure 4**). We measured the expression of genes related to calcium handling, contractile mechanism, autophagy, ER stress, metabolism, and sarcomere structure. Our results showed many contractile functions and calcium handling relevant genes were up regulated in both MM treated WT and PLN R9C KI cells, such as PLN and CASQ2 (**Figure 4A** and **4B**). Notably, the KI cells showed significantly increased MYH6 and decreased expression of MYH7 compared to WT cells, and the MYH7/MYH6 ratio (an indicator of maturation of cardiomyocyte) was greatly reduced in mutant cells both before and after MM treatment (**Supplementary Figure 4A** to **4C**). Similarly, the SNC5A gene, which encodes sodium channel NaV1.5, was also decreased in mutant cells compared to WT (**Supplementary Figure 4D**). We also observed the key proteins of sarcomere and mitochondria function, such as MYL2 and MFN2, were only upregulated in WT cell after MM treatment (**Figure 4C**, **supplementary Figure 4E**). Instead, many other genes related to apoptosis, protein degradation, ER stress, and autophagy, such as BCL2, CALR, BAG1/3, HSPA1A, HSPA5, HSP90B1, and LAMP2., were highly upregulated in MM treated mutant cells, while minor or no change was observed in WT (**Figure 4D** to **4H, supplementary Figure 4F** to **4H**). Overall, the mRNA expression profile of iPSC-CMs from both groups suggested MM treatment promoted functional maturation in WT cells, while increased proteostasis stress in PLN-R9C KI cells.

**Figure 4.**
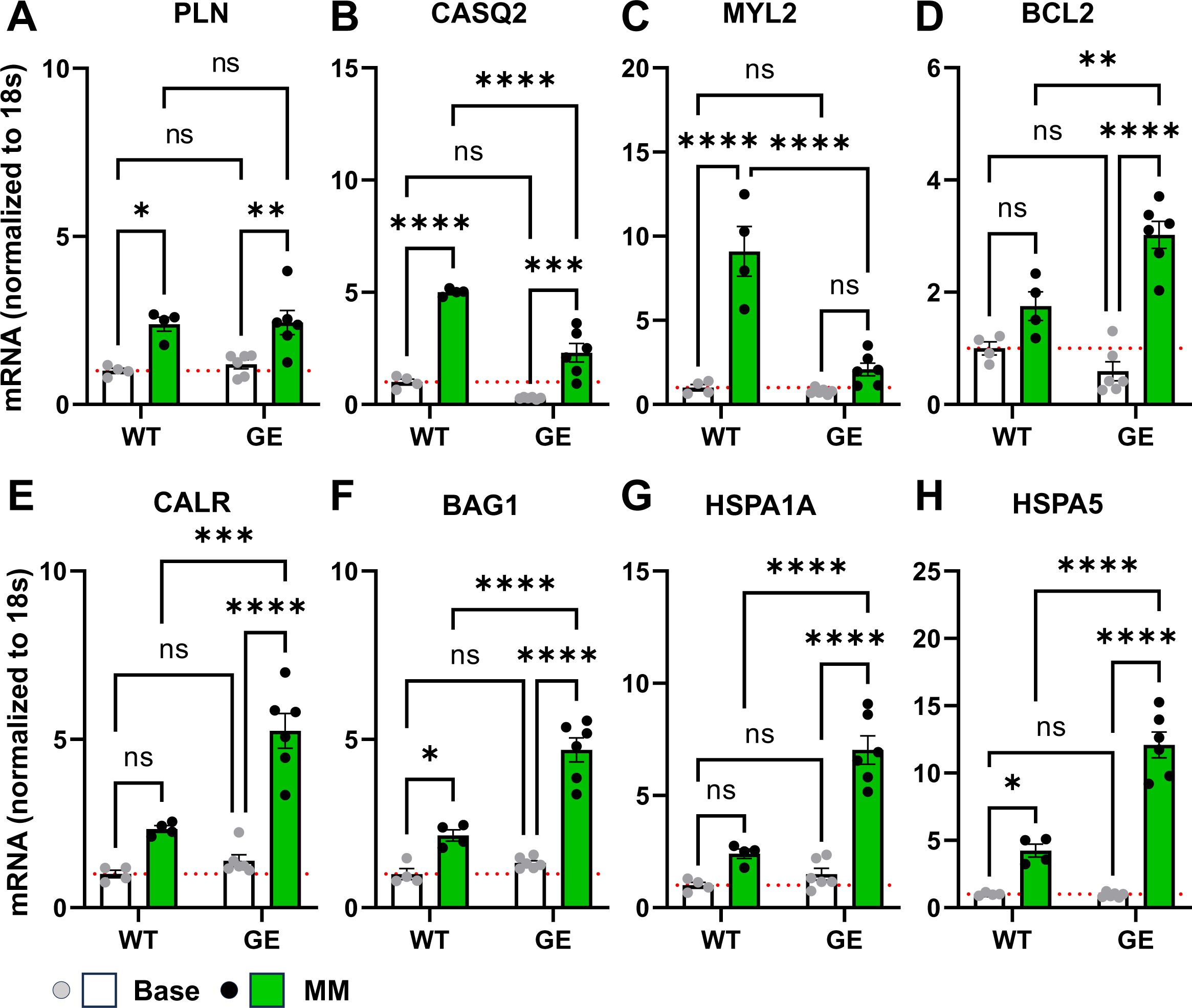
Validation of MM treated WT and R9C iPSC-CMs RNA-seq data with qPCR. A and B. Increased expression of calcium handling relevant genes PLN (A) and CASQ2 (B) in both WT and GE iPSC-CMs after MM treatment. C. Increased expression of sarcomere protein MYL2 specifically in WT iPSC-CMs after MM treatment. D to H. The expression of apoptosis gene BCL2 (D), ER stress marker CALR (E), and protein homeostasis relevant genes BAG1 (F), HSPA1A (G) and HSPA5 (H) were significantly upregulated in PLN R9C KI iPSC-CMs after MM treatment, while only mild or no increases are observed in the WT group. Results from at least 4 independent experiments. NS, no significance, *p<0.05, **p<0.01, ***p<0.001, and ****p<0.0001 by 2-way ANOVA test followed by Holm-Sidak method.

### Interrupted proteostasis and autophagy flux in PLN R9C KI iPSC-CMs upon maturation challenge

To further validate the expression profile of WT and KI cells, we used western blot to quantify the abundance of key proteins in WT and mutant cells before and after MM treatment. Previous studies have shown that, the PLN R9C mutation affects the dynamics of the formation of PLN pentamers as well as the PLN-SERCA2a complex, which affects PLN phosphorylation (cardiac functional regulation) as well as PLN ubiquitination (proteostasis).^26^ As proteostasis is key to the maintenance of cardiac function, we also examined the PLN level and polymerization. Our result shows the PLN protein level in R9C mutant cells is greatly reduced compared to WT, potentially due to the activation of nonsense-mediated RNA decay (NMD) pathway and decreased binding of PLN to SERCA2a (**Figure 5A**).^26, 27^ After being challenged by MM, the PLN expression level was increased by ∼33.5% in WT group, while the PLN level in R9C cells were significantly upregulated by ∼72.4%, indicating an increased demand of dynamic functional regulation in more mature cardiomyocytes (**Figure 5B**). Notably, the pentamer/monomer ratio of PLN was significantly increased in R9C cells, but not in WT cell, indicating more PLN forms pentamers in mutant cells, likely due to the insufficient degradation of excess PLN protein (**Figure 5C**). As autophagy is essential for PLN degradation in lysosomes, we next examined the autophagic flux in both WT and PLN R9C cells by measurement of the LC-3 II/LC-3 I ratio (**Figure 5D**).^28, 29^ Interestingly, while WT and PLN R9C cells exhibited unchanged autophagic fluxes at base line, MM treatment has increased the autophagic stress specifically in mutant cells, as the LC-3 II/LC-3 I ratios were up-regulated by 187.6% for PLN R9C MM cells compared to their WT control (**Figure 5E**). In accordance with the increased autophagic stress in mutant cells, quantification of HSP70, a master regulator of protein degradation, is significantly decreased in PLN R9C cells in response to MM treatment (**Figure 5D** and **5F**). Increased autophagy and protein homeostasis stress is known to promote the mis-regulation of cellular ROS and mitochondrial function. We used a ROS indicator and mitochondrial trackers to evaluate the consequences of MM challenge in both WT and mutant cells (**Supplementary Figure 5A**). Our MM treatment results showed both cellular and mitochondrial ROS levels were increased by 81.2% in mutant cells, while no change was detected in WT group (**Supplementary Figure 5B**).

**Figure 5.**
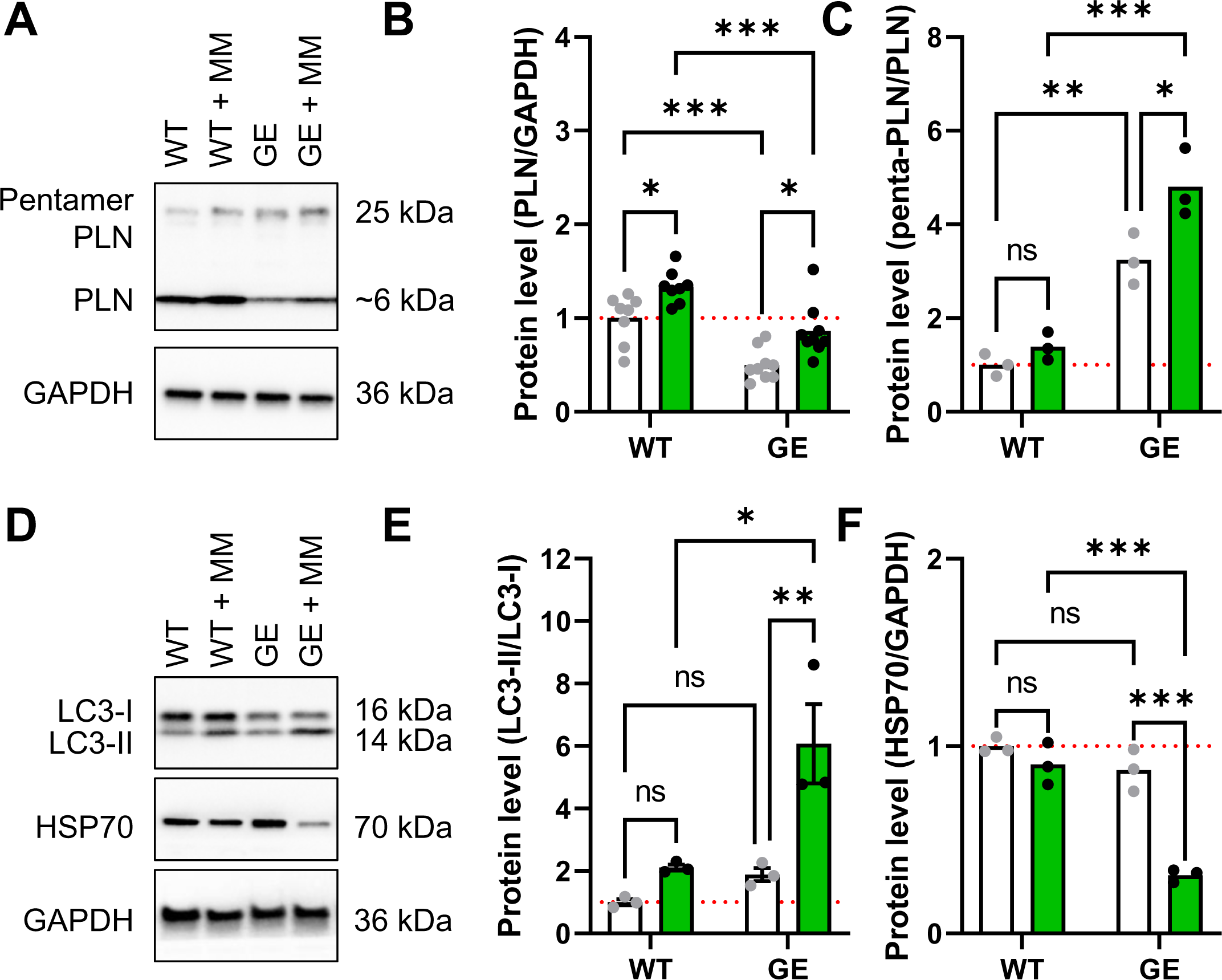
MM increased protein aggregation in PLN R9CKI iPSC-CMs. A. Western blot quantification of monomer and pentamer PLN abundance in WT and PLN R9C KI iPSC-CMs. B. Quantification showed significantly decreased PLN protein in GE iPSC-CMs compared to WT cells both before and after MM treatment, although MM enhanced the PLN level in both groups. Results from at least 7 independent experiments. C. The PLN aggregation, as evidenced by the pentamer/monomer PLN ratio was unchanged in MM treated WT cells, while was remarkably higher in the GE group, both before and after MM treatment. Results from at least 3 independent experiments. D. Western blot analysis of the of LC3-I, LC3-II, and HSP70 expression. E. Autophagy is unchanged in WT cells before and after the functional stress by MM, while compromised autophagic flux is evidenced by the significantly increased LC3-II/I ratio in GE iPSC-CMs after MM treatment. Results from at least 3 independent experiments. F. HSP70 protein level is unchanged in GE iPSC-CM and WT cells at base level, but significantly decreased in MM treated GE iPSC-CMs compared to WT group. Results from at least 3 independent experiments. NS, no significance, *p<0.05, **p<0.01, and ***p<0.001 by 2-way ANOVA test followed by Holm-Sidak method.

### Autophagic dysfunction in DCM patient specific iPSC-CMs

To validate our findings of the pathological mechanism of PLN R9C in human samples. We recruited a male DCM patient carrying PLN R9C mutation, who had an early on-set DCM in his 40s. Patient fibroblast samples were collected and reprogrammed to iPSCs using standard protocols (see methods). The patient-specific iPSC clones were characterized by pluripotency staining and in-vitro 3-germ layer differentiation methods. (**Supplemental Figure 6A** to **6C**) The PLN R9C heterozygous mutation of iPSCs were confirmed by genotyping, and the iPSCs passaged 20 times before cardiomyocyte differentiation and further functional measurements. (**Supplemental Figure 6D** to **6F**) Like our observations in PLN R9C heterozygous KI iPSC-CMs, DCM patient-specific (PA) iPSC-CMs exhibited near-normal morphology and function at base level. (**Figure 6A**) Yet the β-adrenergic signaling in PA cells was impaired compared to WT cells, as evidenced by unchanged transient rise time and calcium recycling rate (**Figure 6B** and **6C**), mild increase of beating rate (**Figure 6D**), and unchanged contraction/relaxation function in response to ISO treatment. Also, the time course of the contraction, such as contractile duration, relaxation duration, and full duration of PA cells show much smaller changes upon ISO challenge compared to WT. (**Supplemental Figure 6G** to **6I**) All these observations suggested that, like GE cells, the PLN R9C protein has blunted β-adrenergic signaling in PA iPSC-CMs. We also applied functional challenges to the PA iPSC-CMs by MM treatment. (**Figure 7A**) Similar to our findings in PLN R9C KI cells, MM treatment has impaired the calcium handling and contractile function of PA iPSC-CMs in terms of decreased calcium transient amplitude, slower calcium recycling, and decreased contractile/relaxation velocity (**Figure 7B** to **7E**). Accordingly, we also observed overloaded diastolic calcium, and prolonged contraction/relaxation duration in PA iPSC-CMs after MM treatment (**Supplementary Figure 7A** to **7C**).

**Figure 6:**
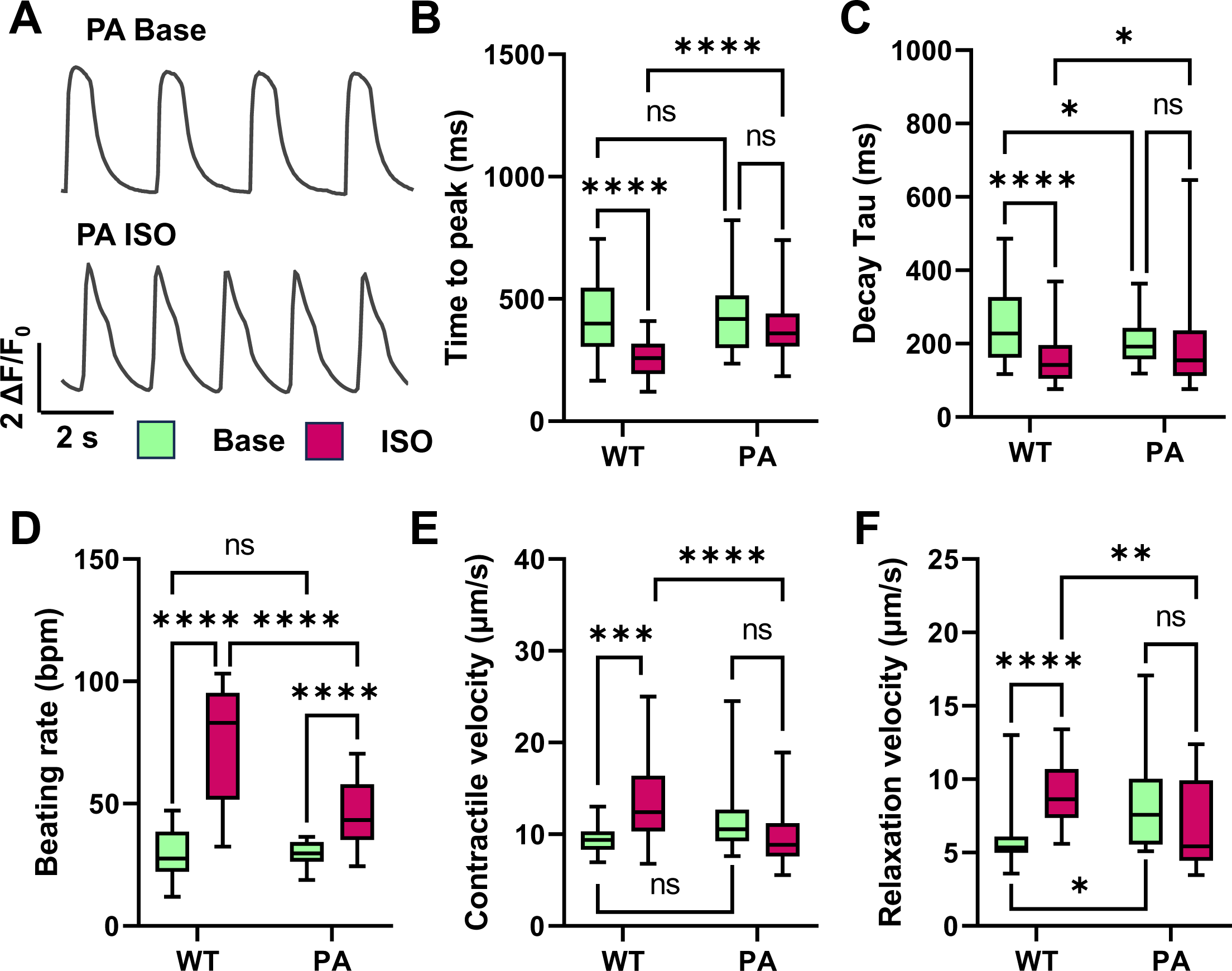
Functional profiling of patient-specific PLN R9C iPSC-CMs. A. Representative Fluo-4 calcium transient trace in patient-specific (PA) iPSC-CMs before and after ISO treatment. B to D. β-adrenergic activation induced no or milder changes in calcium handling capacity, such as transient rise time (B), transient decay Tau (C), and beating rate (D) in PA iPSC-CMs compared to WT cells. N > 101 cells of WT, PA, WT ISO, and PA ISO groups from at least 3 independent experiments. E and F. Traction force microscopy analysis showed slightly increased contraction (E) and relaxation velocity (F) in PA iPSC-CMs at base level compared to WT cells. Yet contractile function of PA iPSC-CMs is significantly reduced compared to WT cells after ISO treatment, as no functional improvement is induced by ISO in PA cells. N > 26 ROIs of WT, PA, WT ISO, and PA ISO groups from at least 3 independent experiments. NS, no significance, *p<0.05, ***p<0.001, and ****p<0.0001 by 2-way ANOVA test followed by Holm-Sidak method.

**Figure 7:**
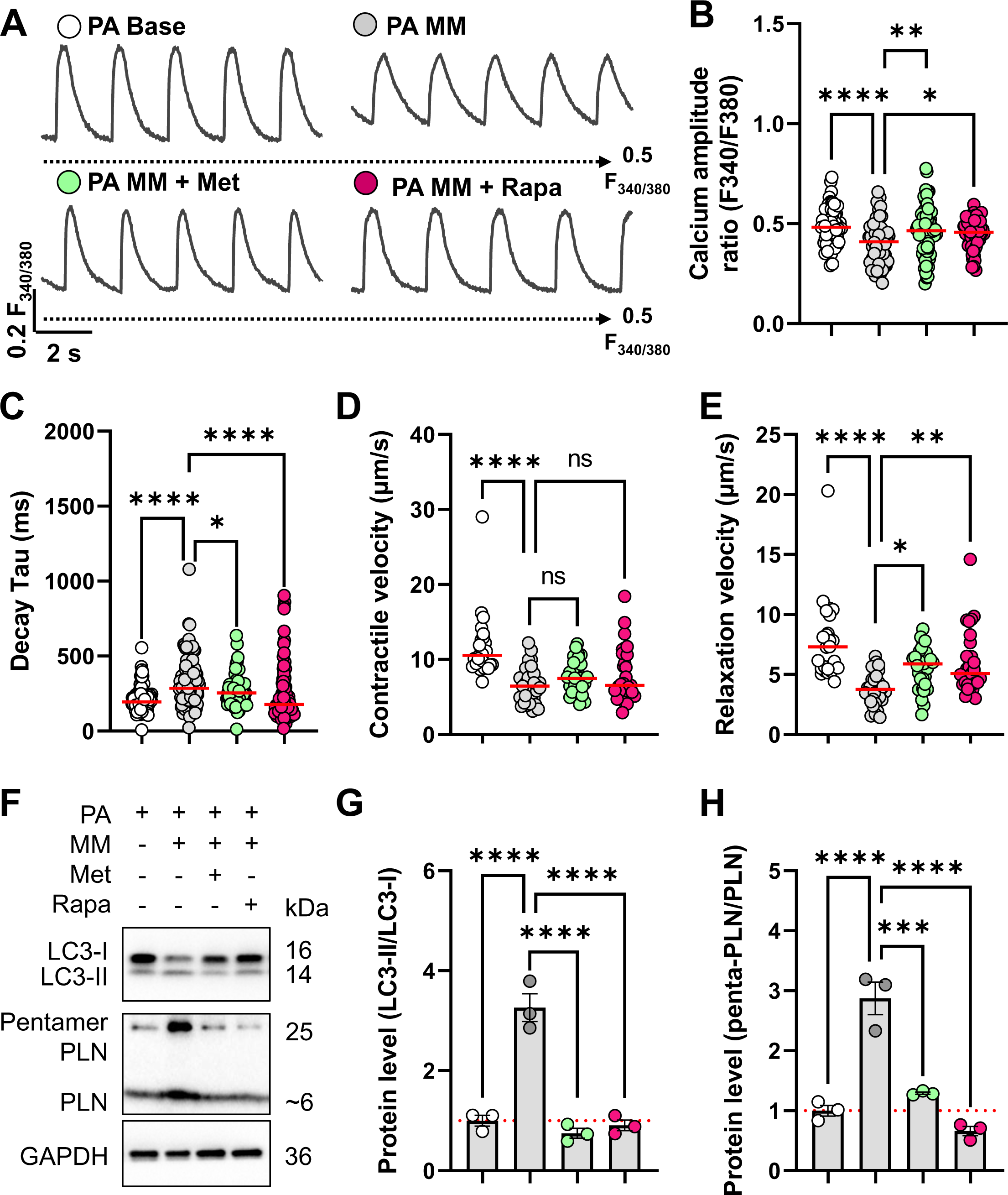
Activation of autophagy restored cardiac function of patient-specific PLN R9C iPSC-CMs. A. Representative ratiometric Fura-2 AM calcium transient trace in PA iPSC-CMs at baseline, after MM treatment, and MM treatment in presence of 2 mM metformin or 0.5 µM rapamycin. Black dashed line indicates the F340/F380 ratio of 0.5. B to C. Long-term MM treatment enhanced Calcium transient amplitude in WT cells, but not in the GE cells. N > 78 ROIs of WT, GE, WT MM, and GE MM groups from at least 3 independent experiments. D and E. TFM measurement of contraction (D) and relaxation (E) velocity in WT and PA cells after MM treatment. N > 26 ROIs of PA base, PA MM, PA MM with metformin, and PA MM with rapamycin groups from at least 3 independent experiments. NS, no significance, *p<0.05, **p<0.01, and ****p<0.0001 by 1-way ANOVA test followed by Holm-Sidak method. F. Western blot quantification of the abundance of LC3-II, LC3-I, as well as monomer and pentamer PLN in PA base, PA MM, PA MM with metformin, and PA MM with rapamycin groups. G and H. Quantification of western blot showed MM treatment has increased the autophagic stress (LC3-II/LC3-I) level and promote pentamer aggregation of PLN (penta-PLN/mono-PLN), which were mitigated by the activation of autophagy by metformin and rapamycin treatment. Results from 3 independent experiments. *p<0.05, ***p<0.001, and ****p<0.0001 by 1-way ANOVA test followed by Holm-Sidak method.

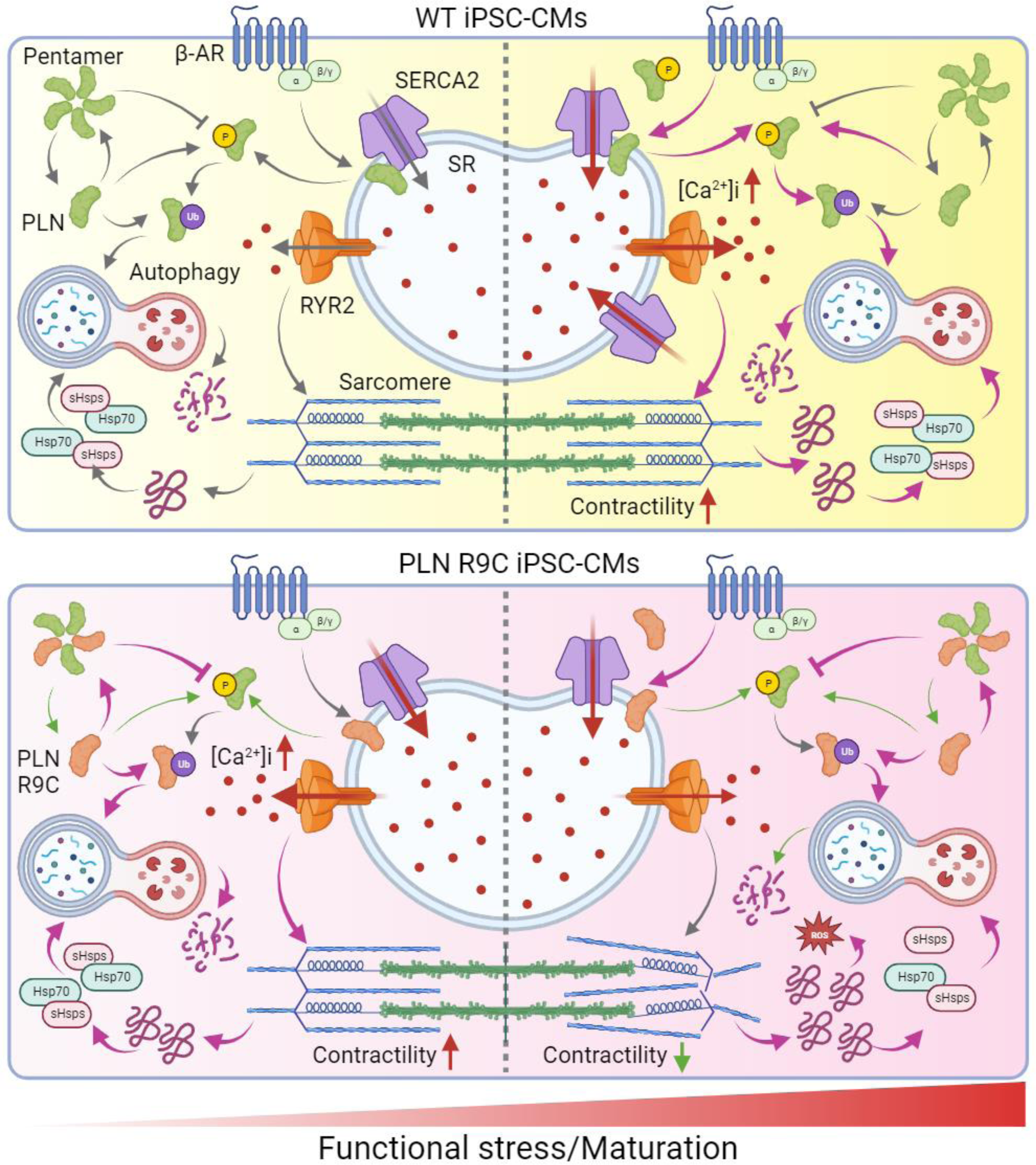

### Activation of autophagy restored proteostasis and cardiac function in patient-specific iPSC-CMs

As both our RNA-seq results and functional study highlighted the mis-regulation of protein homeostasis as a key signaling pathway during the functional stress-induced pathogenesis in PLN R9C cardiac cells, we sought to alleviate the functional deficiency of diseased cells through activation of autophagic signaling pathways. In our study, we used both metformin and rapamycin to treat patient-specific iPSC-CMs, which are known to promote autophagy signaling through the activation of AMPK or the inhibition of mTOR.^30, 31^ In our observation, treatment with 2 mM metformin (Met) or 0.5 uM rapamycin (Rapa) for 7 days during MM challenge at least partially restored the function of iPSC-CMs (**Figure 7A**). Both calcium imaging and traction force microscopy analysis confirmed that activation of autophagy enhanced calcium transient amplitude, accelerated calcium recycling, restored diastolic calcium level, and improved contraction and relaxation functions in functionally stressed PA cells (**Figure 7B** to **7E, Supplementary Figure 7A** to **7C**). In addition, morphological measurements of sarcomere arrangement by immunostaining also confirmed, the activation of autophagy flux was able to maintain the structural integrity of the contractile mechanism in diseased cardiomyocytes under stress, as both Metformin and Rapamycin treatment have improved the arrangement of sarcomere structures (**Supplementary Figure 7D** and **7E**). Further analysis by western blot confirmed the activation of autophagic flux by Metformin and Rapamycin, as the LC3II/LC3I ratio was significantly reduced in the treated PA cells (**Figure 7F** and **7G**). As a result, the PLN protein level and PLN aggregation (measured as pentamer PLN/Monomer PLN ratio) were both reduced after Metformin and Rapamycin treatment compared to the MM group (**Figure 7F** and **7H**). As the activation of autophagy ameliorated the protein homeostasis and ER stress, we also observed a reduction of ROS levels in DCM iPSC-CMs in metformin and rapamycin treatment groups (**Figure 7F**). Together, all the evidence suggests that the activation of autophagy signaling has partially rescued the morphological and functional deficiencies in DCM patient iPSC-CMs with PLN R9C mutation, likely through restoration of protein homeostasis in the cardiomyocytes.

## Discussion

So far, multiple PLN mutations and genetic variants have been reported in cardiomyopathy and heart failure patients, which include: 1. missense mutations such as p.Arg9Cys (R9C), p.Arg9His (R9H), and p.Arg25Cys (R25C); 2. nonsense mutations such as p.Leu39Ter (L39X) and p.Glu2Ter (E2X), 3. deletion mutation p.Arg14del (R14del), as well as other genetic variants.^14^ Among all the PLN mutations, R14del is perhaps the most well-known one, which has been studied and characterized in many animal and cell line models. Our current understanding of the mutation is PLN-R14del has increased association with SERCA2a, which leads to increased SERCA2a inhibition and decreased SR Calcium uptake and depressed contractility. As PLN-R-14 lacks the phosphorylation at Ser-16, the inhibition of the SERCA2a becomes chronic and cannot be relieved by β-adrenergic stimulation.^32^ The decreased SR calcium transport velocity leads to the overall increase of diastolic calcium level and promotes the arrhythmogenesis in PLN R14del patients.^12, 33, 34^ Similar to PLN-R14del, a previous study showed PLN-R25C mutation also exhibited enhanced interaction with SERCA2a, thus lead to super-inhibition of SERCA2a, decreased SR calcium load and calcium transient, and impaired contractile function. PLN-R25C could be phosphorylated by PKA, which prevents the inhibition of SERCA2a by PLN.^7^ Thus β-adrenergic stimulation can partially relieve the functional deficiency in PLN R25C cardiomyocytes. However, excess activation of β-adrenergic signaling induces increased CaMKII activity and hyper-phosphorylation of RyR2-2814, thus promoting arrhythmogenesis in PLN R25C patients under stressed conditions. PLN R9C was known to lead to early on-set severe DCM without an arrhythmia phenotype.^7^ Unlike the enhanced inhibitory effect on SERCA2a observed in the R14del and R25C mutations, the PLN R9C variant results in nearly complete loss of binding to and inhibition of SERCA2a, underscoring its distinctive role in pathogenesis. Nevertheless, our current understanding of the intricate molecular mechanisms underlying the pathogenesis of PLN R9C remains incomplete.

In our current study, by establishing the isogeneic R9C knock-in iPSC line, we systematically investigated the role of PLN R9C mutation in the context of human cardiomyocytes. Our results partially agree with previous iPSC modeling study in that the β-adrenergic signaling was attenuated in the KI iPSC-CMs.^20^ However, in our functional assays, the PLN R9C KI iPSC-CMs showed near-normal or even enhanced function at baseline in many aspects. For example, the PLN R9C iPSC-CMs exhibited unchanged beating rate and calcium transient amplitude, while the calcium recycling rate (Decay Tau) and relaxation velocity were increased in mutant cells. This is in line with previous reports that R9C mutation has significantly decreased the interaction between PLN and SERCA2a, thus enhancing the SERCA activity at basal level. Indeed, our Fura-2 based calcium imaging also showed a decreased diastolic calcium level in R9C cells compared to WT at baseline, confirming the enhanced calcium recycling in mutant cells. (**Supplemental Figure 2D**) On the other hand, one of the most significant functional deficiencies we observed in R9C iPSC-CMs is the blunted response to β-adrenergic signaling, as the same ISO challenge only results in milder functional gain in mutant cells, as shown in calcium recycling rate, beating rate, as well as the contractile and relaxation durations. Indeed, previous studies suggested that the conformational change of PLN-R9C mutation reduced the binding and phosphorylation by activated PKA, which reduced the percentage of phosphorylated PLN molecule by 60%.^23^ Moreover, the PLN-R9C-PKA complex has a much slower dissociation rate compared to WT PLN-PKA complex, which suggests the PLN-R9C acts as a trapper of activated PKA, thus further diminishing the β-adrenergic signaling in mutant cardiomyocytes. This is further confirmed by our quantification of the p.Ser16-PLN levels before and after ISO treatment in both mutant and WT iPSC-CMs. Interestingly, the functional performance of the PLN R9C iPSC-CMs, although slightly increased at base level compared to WT cells, is much decreased compared to the WT cells in response to the β-adrenergic challenge, which may indicate a gap between the contractility and functional demand in PLN R9C cardiomyocytes under stressed conditions.

To evaluate how mutant and WT cells respond to functional stress, we applied maturation medium (MM), which promotes the functional maturation of iPSC-CMs in culture.^24, 35^ MM treatment induced the upregulation of genes related to contractile function and calcium handling (pathways highlighted in RNA-seq), such as MYL2 and CASQ2, and enhanced the contractility and calcium handling in WT cells. While at the same time, MM increased stress on protein homeostasis and <u>exacerbated the</u> disease phenotypes in PLN R9C iPSC-CMs.^25^ Indeed, our RNA-seq and GO-term analysis demonstrated significantly increased enrichment of protein homeostasis pathways, such as protein degradation and absorption, in MM-treated PLN R9C cells, but not WT cells, which is plausible when taking into account the elevated contractility of these cells under normal conditions and their inability to adapt functional output in response to demand (due to diminished β-adrenergic signaling). Moreover, the presence of functional challenge also led to impaired autophagy flux, increased cytosolic and mitochondrial ROS, as well as mitochondrial stresses. Notably, while there is almost no change in mRNA level, the protein level of PLN was greatly decreased in PLN R9C cells compared to WT cells, which indicates mutant PLN is likely to be degraded by the protein quality control system at base level. However, after MM treatment, PLN levels were greatly upregulated in mutant cells, suggesting less mutant PLN was degraded through autophagy. As a result of the accumulation of mutant PLNs, their resistance to phosphorylation/degradation, as well as the increased oxidative stress, PLNs in mutant cells are more likely to form pentamers in the MM group compared to baseline.^19^ A deficiency in the protein quality control system could limit the renewal of key structural related proteins, which led to the disarrangement of the myofilaments in mutant cells under prolonged stress.

All these findings suggest that the PLN R9C mutation may contribute to early on-set dilated cardiomyopathies through the following two mechanisms: 1: PLN R9C has reduced binding and phosphorylation by PKA, which means the mutant cells are not sensitive to β-adrenergic signaling. Thus, although the cells behave normally at basal level, their gain of function after β-adrenergic challenge is insufficient to satisfy the increased functional demand in a stressed heart. As a result, our body will compensate the functional deficiency with a higher amount of adrenaline signaling, which further increases the cardiac function in the short term but will also lead to long-term structural and functional remodeling of the heart, which include the loss of cardiomyocyte, increased fibrosis, and heart failure.^36, 37^ 2: PLN R9C exhibits a nearly complete loss of its inhibitory effect on SERCA2a, resulting in elevated contractile function of the heart at baseline. This poses an intensified challenge to protein quality control. Thus, in response to functionally challenged conditions, such as MM treatment, the mutant cells will endure an increased stress in protein homeostasis, as evidenced by our RNA-seq data and autophagy flux measurement. Increased misfolded proteins also lead to ER stress, ROS overload, and mitochondrial dysfunction, which are also key mechanisms underlying the pathological role of PLN R9C mutation.

Autophagy is a highly conserved process through which long-lived/damaged proteins and organelles are degraded within lysosomes, which keeps live cells in homeostasis.^38^ Although autophagic activity is low in cardiomyocytes at a basal level, it can be activated and serve as a protective mechanism against toxic protein aggregation induced by cardiac stress (such as energy restriction or high function demand).^39^ Autophagy plays an important role in the pathogenesis of inherited cardiomyopathies, as they are caused by mutant/misfolded protein. Indeed, defects in autophagy have been identified in a wide range of cardiac diseases thus far, which include DCM, hypertrophic cardiomyopathy (HCM), diabetic cardiomyopathy, desmin-related cardiomyopathy (DRM), and Danon disease.^40^ Due to the critical role of autophagy in cardiomyopathy, the therapeutic benefits of activation of autophagy (either by over expression of autophagy proteins, such as ATG7 or UBC9, or by activation of autophagy signaling pathways by drugs, such as rapamycin, metformin, or resveratrol.) have been tested in mouse disease models.^41–45^ Many studies have reported that the activation of autophagy was able to eliminate protein aggregation, reduce hypertrophy and fibrosis, improve the cardiac systolic and diastolic functions, and extend the life span in the animal disease models of DCM, HCM, and age-related CM.^46–48^ In our current study, we found that PLN R9C iPSC-CMs showed near-normal cardiac function at base level, while the level of PLN is significantly decreased, which may indicate a protein quality control mechanism that reduces the PLN-R9C and compensate the function of cardiomyocytes. However, this mechanism is no longer functional in response to long-term functional challenges, which is evidenced by the activation of protein homeostasis related signaling pathways in RNA-seq. Also, decreased LC3-II/LC3-I ratios indicate the deficiency of autophagic flux in stressed mutant cells. As a result, significant increases of PLN, especially pentamers, as well as functional deficiencies, were observed in PLN R9C iPSC-CMs after MM treatment. Interestingly, an abnormal unfolded protein response and myofilament alternation were reported in recent studies of other PLN mutations as well, which agrees with our findings and may indicate a general mechanism underlying PLN mutation induced signaling remodeling in DCM.^49, 50^ Likewise previous studies showed that the degradation of PLN in cardiomyocytes depends on phosphorylation and autophagy.^26, 51^ To target the PLN R9C induced functional deficiencies in patient-specific iPSC-CMs, we tested the effects of rapamycin and metformin, FDA-approved mTOR catalytic site inhibitor and AMPK activator.^52, 53^ Similar to previous studies, our results show that activation of autophagic signaling reduces PLN accumulation in diseased cells. Here we show, for the first time, that metformin and rapamycin treatment partially restore sarcomere arrangement and cardiac function in functionally challenged patient PLN R9C iPSC-CMs. However, the responsiveness to the β-adrenergic signaling pathway was not rescued by the activation of autophagy signaling, as the PLN level was still significantly lower in treated cells compared to WT cells.

In summary, our current study has identified the mis-regulation of protein homeostasis as a key pathological mechanism of PLN R9C induced cardiomyopathy under functional challenge. We also find that the activation of autophagic signaling can restore autophagic flux, reduce PLN accumulation, prevent structural remodeling, and improve calcium handling and contractile function in functionally stressed patient cells, which can serve as a potential therapeutic approach for these DCM patients.

## Nonstandard Abbreviations and Acronyms

DCM: dilated cardiomyopathy
KI: knock-in
WT: wild type
PLN: phospholamban
CM: cardiomyocytes
iPSC: induced pluripotent stem cell
ISO: isoproterenol
FACS: fluorescence activated cell sorting
TNNT2: cardiac troponin T type 2
MM: maturation medium

## Supplementary Material

Supplementary material is available at XXX Journal online, including:

Detailed Methods

Figures S1–S7

Tables S1–S3

References

## Author’s Contributions

Conceptualization, H.W.; Methodology, H.W., R.J.B., Q.Y., K.S., Q.L., and Y.T.; Formal Analysis, H.W.; Y.S.; Q.L., Investigation, H.W., R.J.B., Q.Y., Q.L.; Resources, Y.T., S.Y.C., J.W.; Data Curation, H.W., Y.S.; Q.L.; Writing—Original Draft Preparation, H.W.; Writing—Review and Editing, R.J.B., S.Y.C., Q.L.; Visualization, H.W., Q.Y, Q.L.; Supervision, H.W. and Q.L.; Project Administration, H.W. and Q.L.; Funding Acquisition, H.W., S.Y.C., and Q.L. All authors have read and agreed to the published version of the manuscript.

## Acknowledgements

We thank Yan Zhuge at Stanford Cardiovascular Institute for her help with iPSC line (NHLBI BhiPSC-CVD 75N9202D00019). We thank Dr. Simon Watkins and Dr. Claudette St. Croix from the Center for Biologic Imaging (University of Pittsburgh) for their technical support. This work benefited from Cytek Aurora CS Spectral Sorter funded by NIH S10OD032265 (PI: Larry Kane).

---

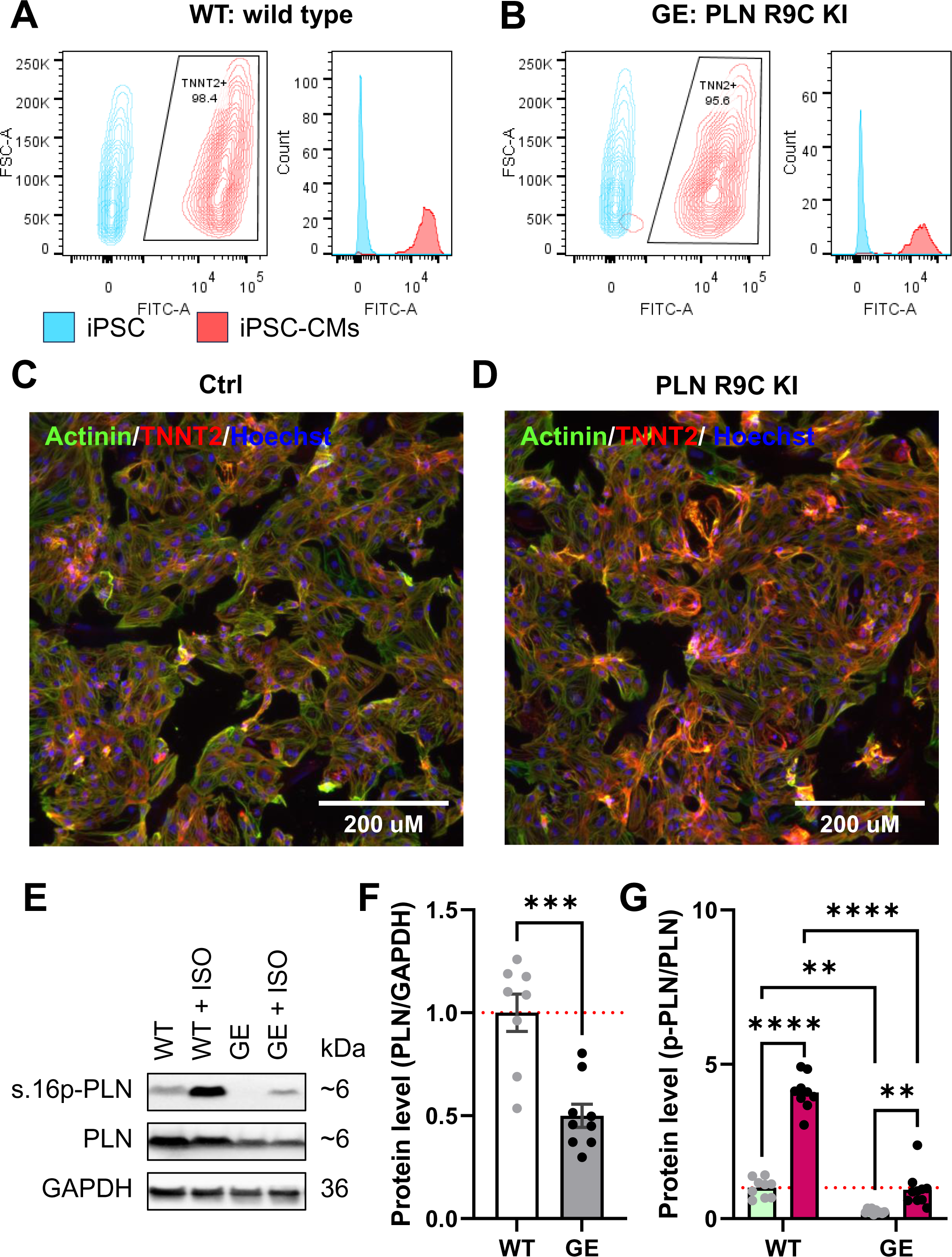

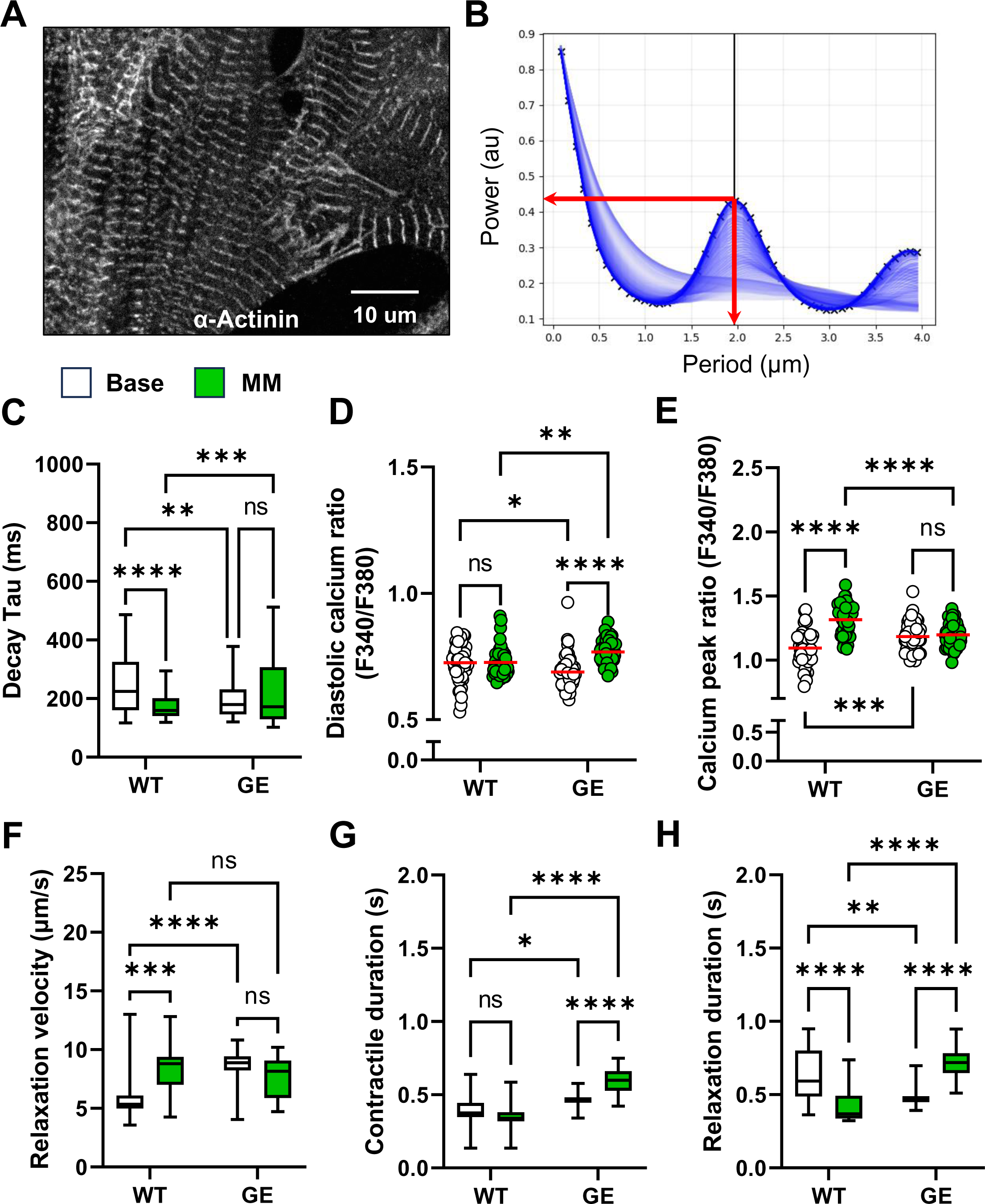

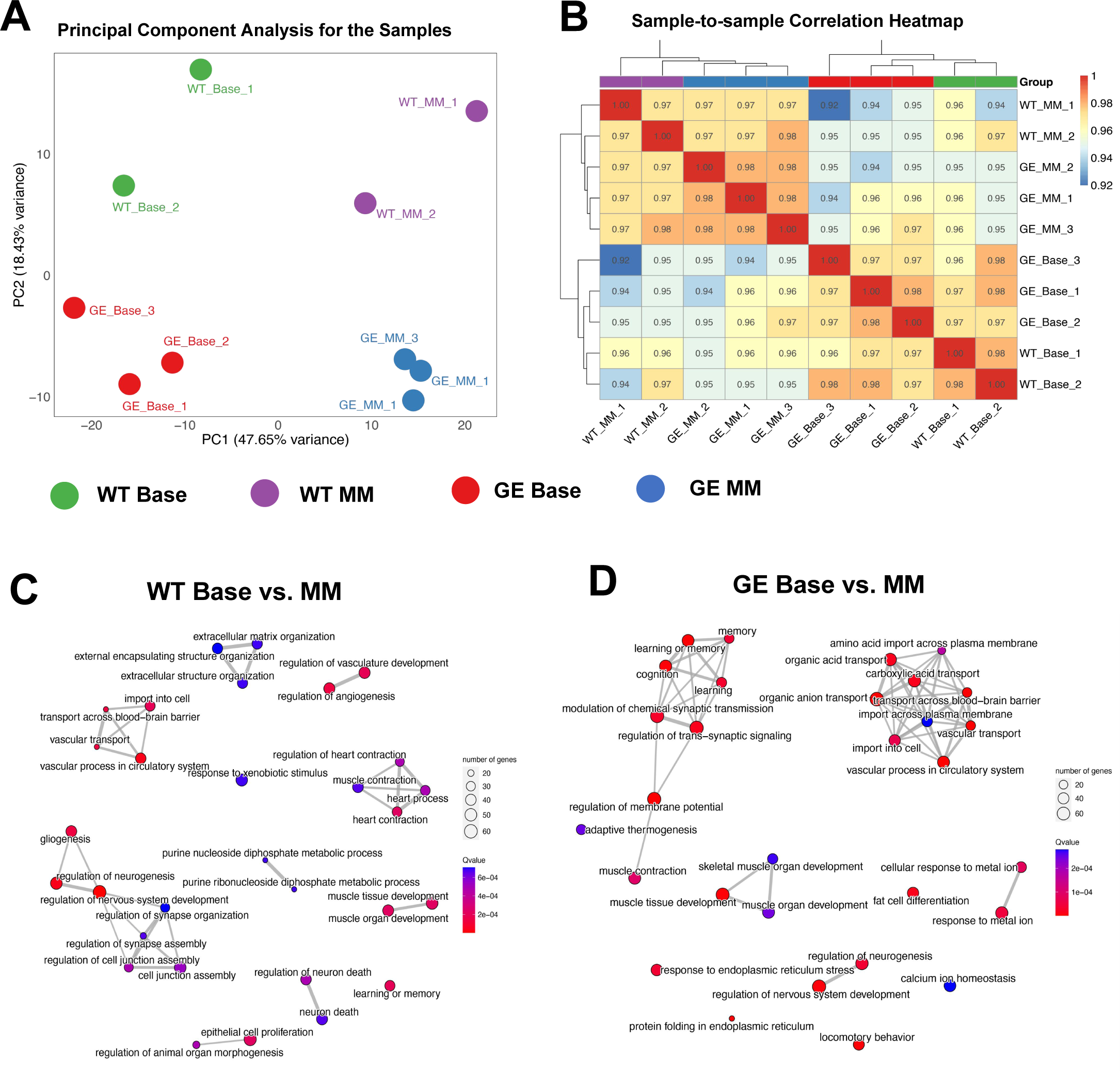

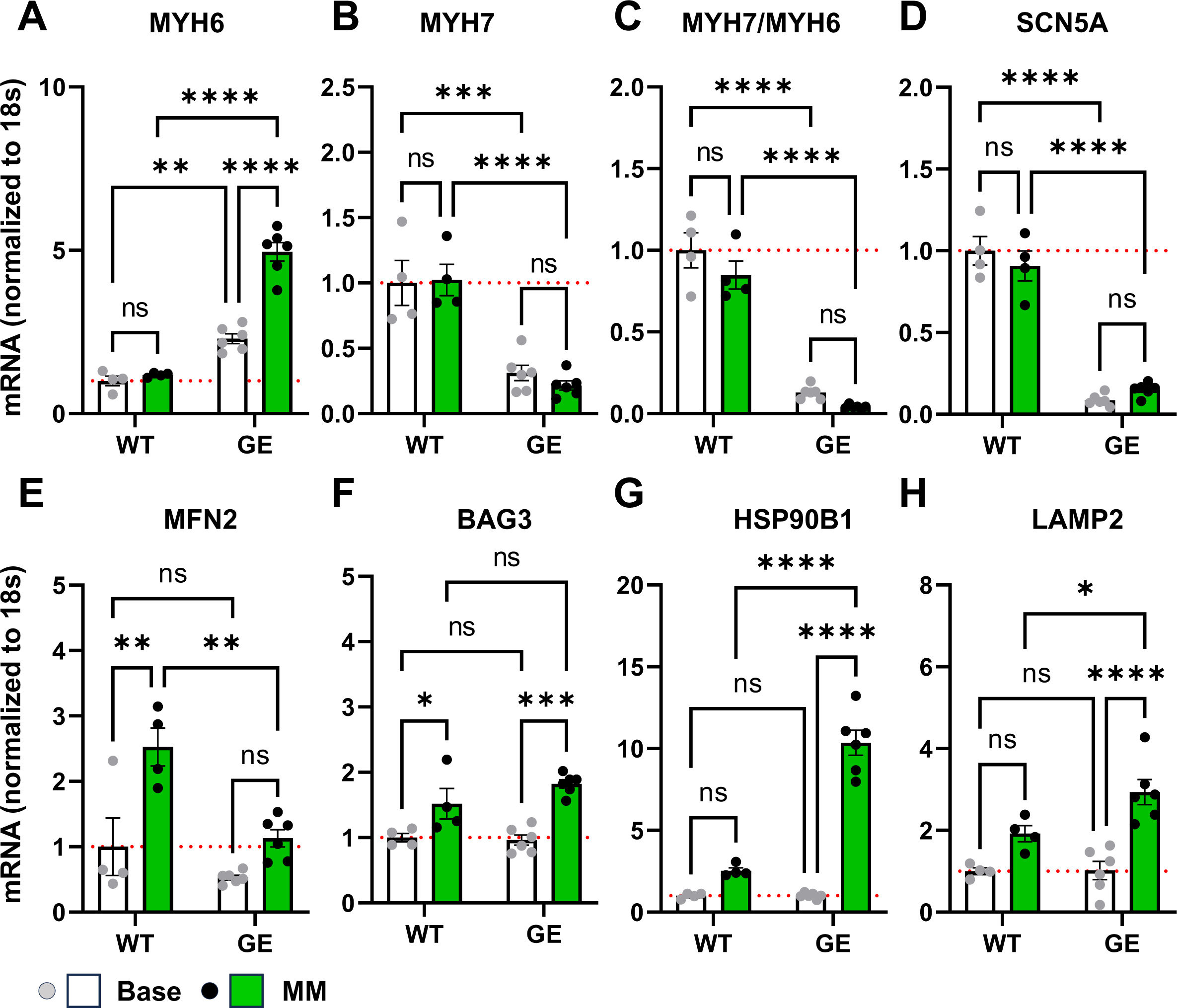

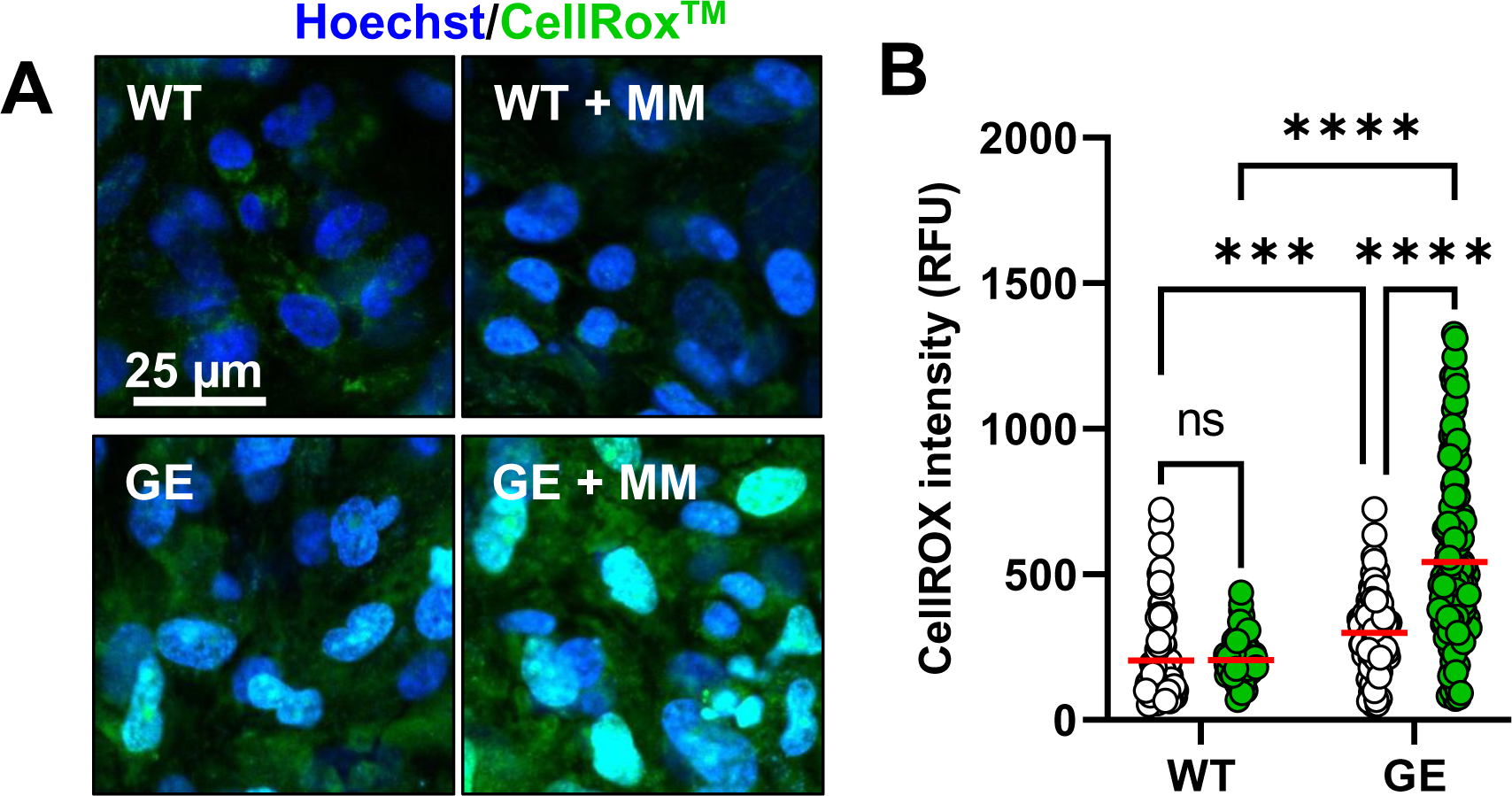

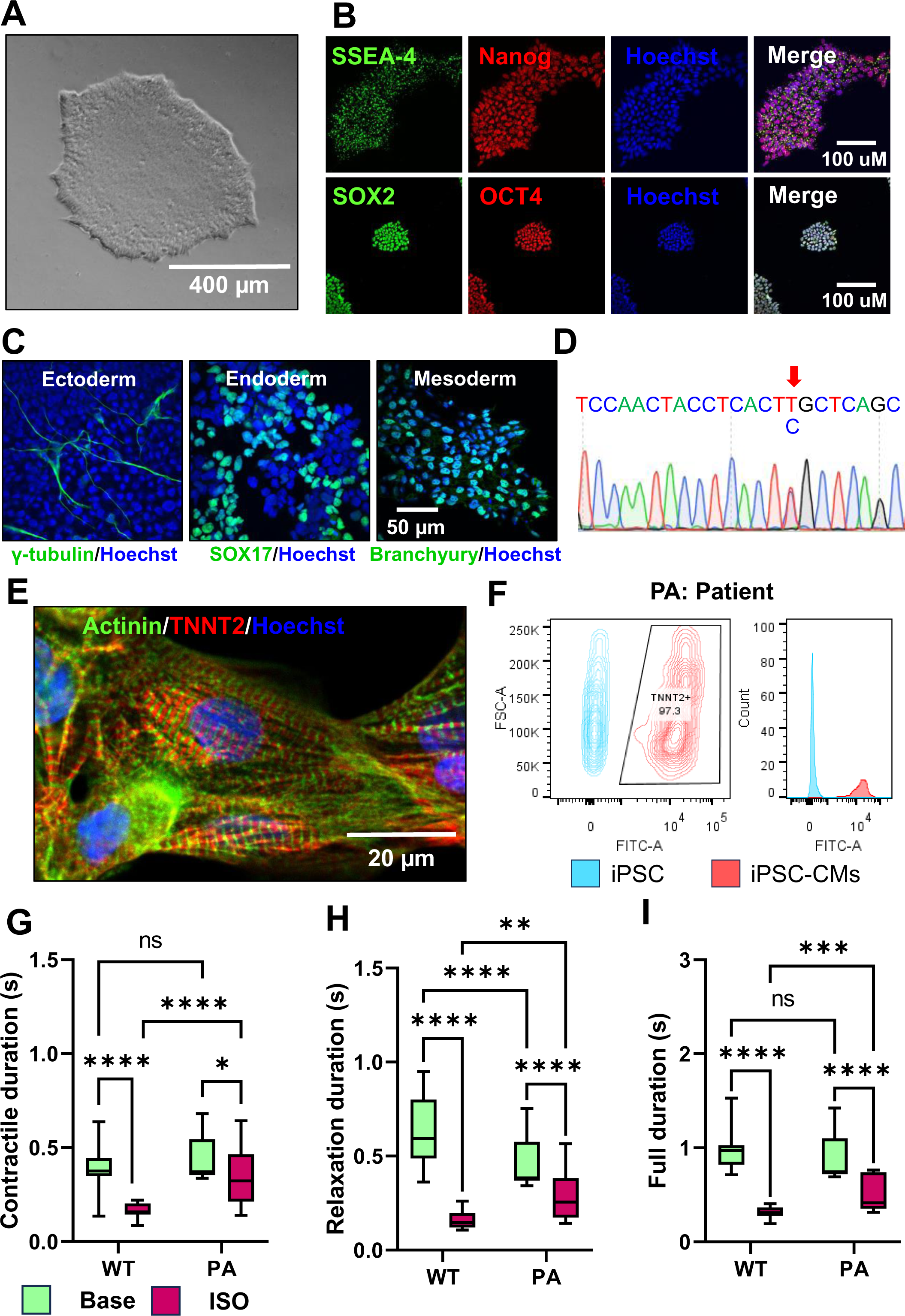

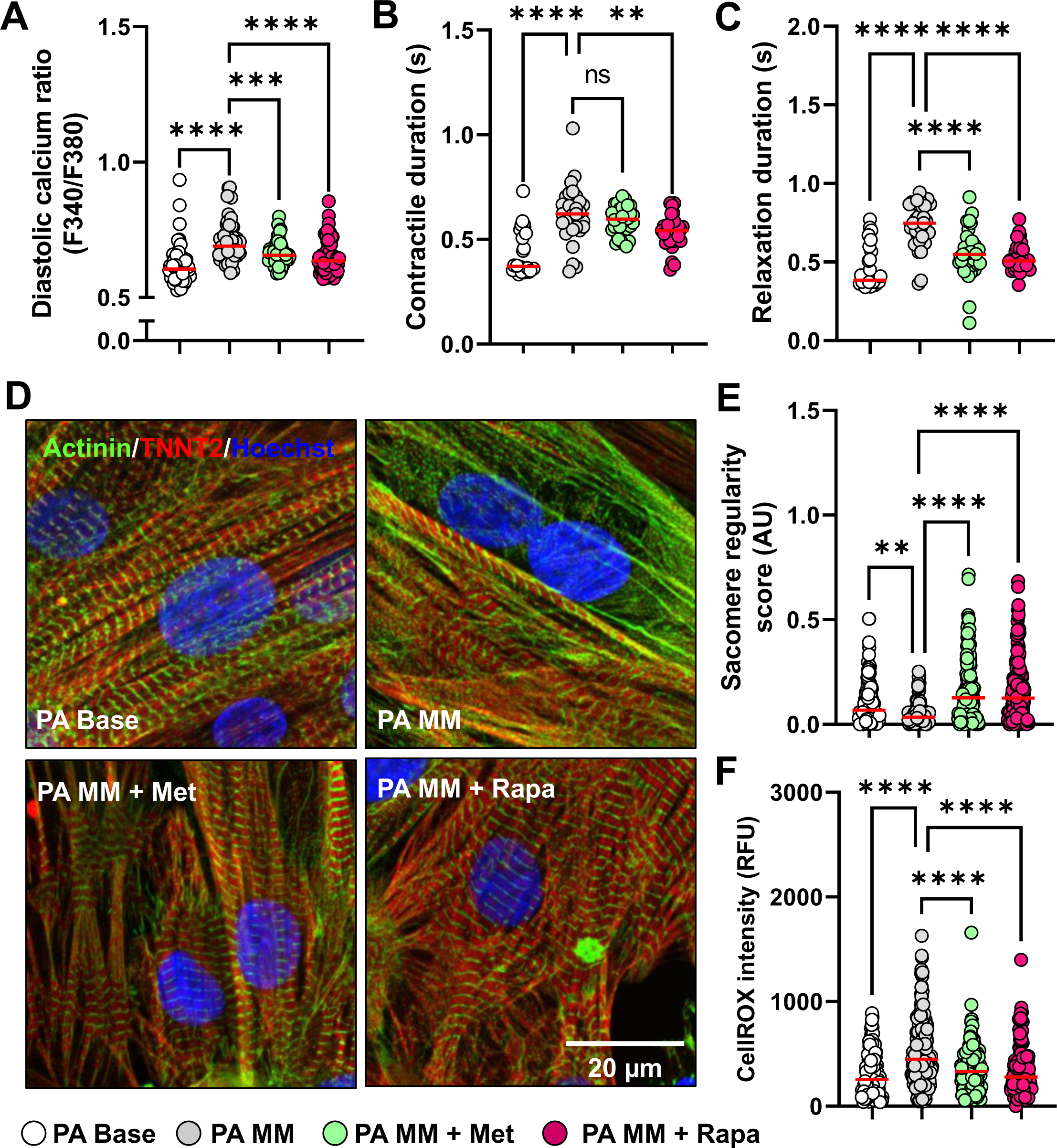

**Supplemental Table 1.**
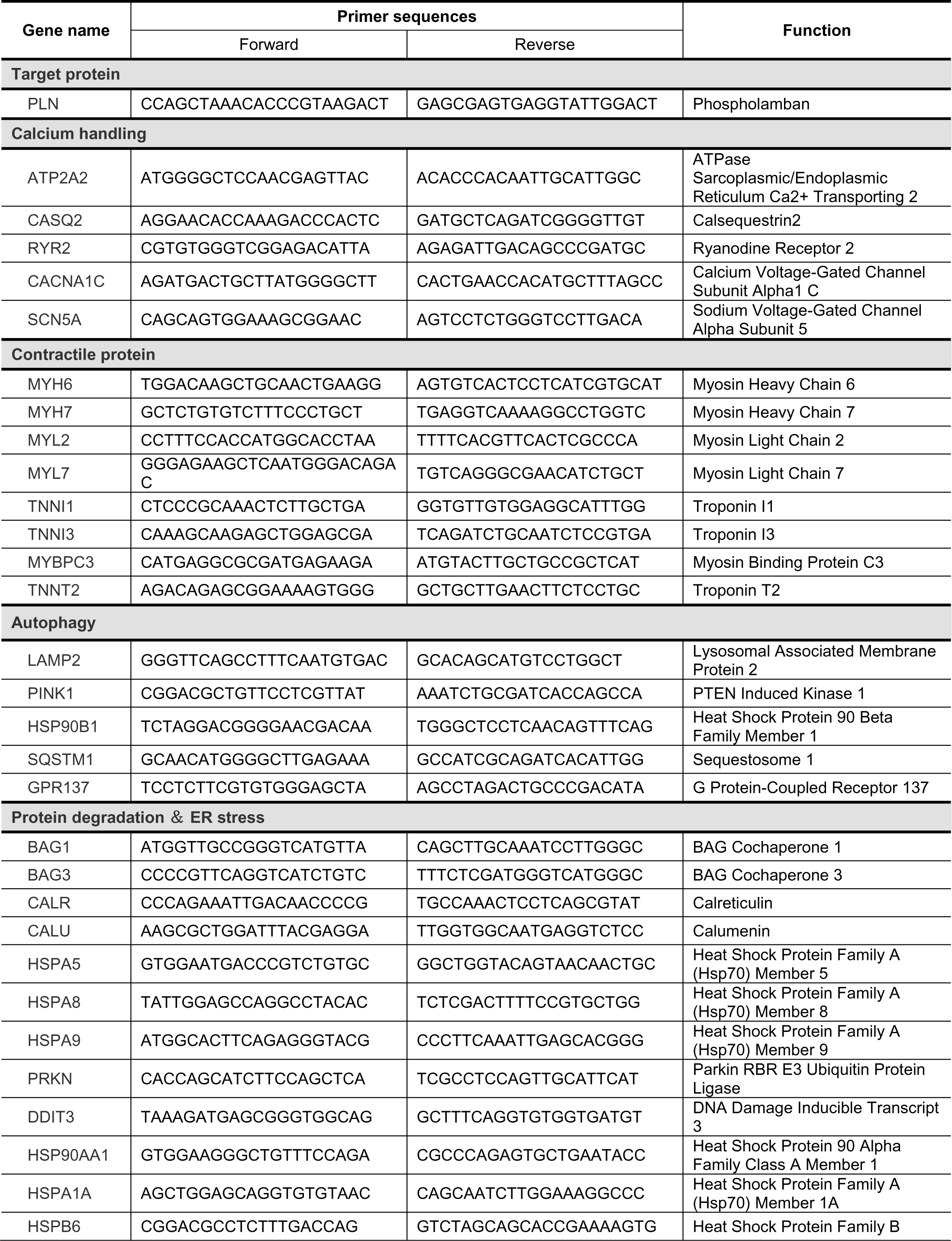
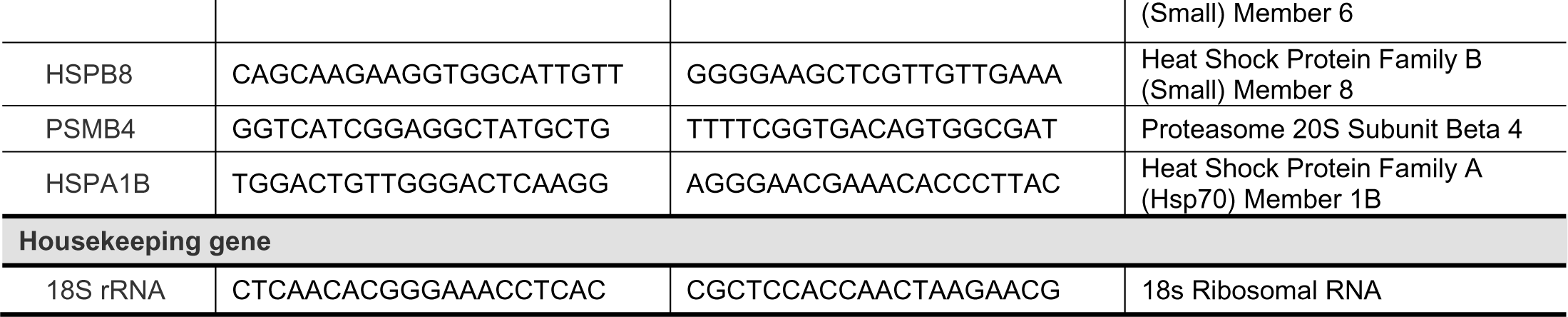
Summary of the sequences of DNA oligos used in the current study.

## References

1. Bers DM. Cardiac excitation–contraction coupling. Nature 2002;415:198–205.

2. Kranias EG, Hajjar RJ. Modulation of Cardiac Contractility by the Phopholamban/SERCA2a Regulatome. Circulation Research 2012;110:1646–1660.

3. Karim CB, Zhang Z, Howard EC, Torgersen KD, Thomas DD. Phosphorylation-dependent conformational switch in spin-labeled phospholamban bound to SERCA. J Mol Biol 2006;358:1032–1040.

4. MacLennan DH, Kranias EG. Phospholamban: a crucial regulator of cardiac contractility. Nature Reviews Molecular Cell Biology 2003;4:566–577.

5. Haghighi K, Kolokathis F, Gramolini AO, Waggoner JR, Pater L, Lynch RA, Fan GC, Tsiapras D, Parekh RR, Dorn GW, 2nd, MacLennan DH, Kremastinos DT, Kranias EG. A mutation in the human phospholamban gene, deleting arginine 14, results in lethal, hereditary cardiomyopathy. Proc Natl Acad Sci U S A 2006;103:1388–1393.

6. Li Z, Chen P, Xu J, Yu B, Li X, Wang DW, Wang DW. A PLN nonsense variant causes severe dilated cardiomyopathy in a novel autosomal recessive inheritance mode. Int J Cardiol 2019;279:122–125.

7. Liu GS, Morales A, Vafiadaki E, Lam CK, Cai WF, Haghighi K, Adly G, Hershberger RE, Kranias EG. A novel human R25C-phospholamban mutation is associated with super-inhibition of calcium cycling and ventricular arrhythmia. Cardiovasc Res 2015;107:164–174.

8. Medin M, Hermida-Prieto M, Monserrat L, Laredo R, Rodriguez-Rey JC, Fernandez X, Castro-Beiras A. Mutational screening of phospholamban gene in hypertrophic and idiopathic dilated cardiomyopathy and functional study of the PLN -42 C>G mutation. Eur J Heart Fail 2007;9:37–43.

9. Schmitt JP, Kamisago M, Asahi M, Li GH, Ahmad F, Mende U, Kranias EG, MacLennan DH, Seidman JG, Seidman CE. Dilated cardiomyopathy and heart failure caused by a mutation in phospholamban. Science 2003;299:1410–1413.

10. Landstrom AP, Adekola BA, Bos JM, Ommen SR, Ackerman MJ. PLN-encoded phospholamban mutation in a large cohort of hypertrophic cardiomyopathy cases: summary of the literature and implications for genetic testing. Am Heart J 2011;161:165–171.

11. Haghighi K, Kolokathis F, Pater L, Lynch RA, Asahi M, Gramolini AO, Fan G-C, Tsiapras D, Hahn HS, Adamopoulos S, Liggett SB, Dorn GW, II, MacLennan DH, Kremastinos DT, Kranias EG. Human phospholamban null results in lethal dilated cardiomyopathy revealing a critical difference between mouse and human. The Journal of Clinical Investigation 2003;111:869–876.

12. van der Zwaag PA, van Rijsingen IA, Asimaki A, Jongbloed JD, van Veldhuisen DJ, Wiesfeld AC, Cox MG, van Lochem LT, de Boer RA, Hofstra RM, Christiaans I, van Spaendonck-Zwarts KY, Lekanne dit Deprez RH, Judge DP, Calkins H, Suurmeijer AJ, Hauer RN, Saffitz JE, Wilde AA, van den Berg MP, van Tintelen JP. Phospholamban R14del mutation in patients diagnosed with dilated cardiomyopathy or arrhythmogenic right ventricular cardiomyopathy: evidence supporting the concept of arrhythmogenic cardiomyopathy. Eur J Heart Fail 2012;14:1199–1207.

13. van Rijsingen IA, van der Zwaag PA, Groeneweg JA, Nannenberg EA, Jongbloed JD, Zwinderman AH, Pinto YM, Dit Deprez RH, Post JG, Tan HL, de Boer RA, Hauer RN, Christiaans I, van den Berg MP, van Tintelen JP, Wilde AA. Outcome in phospholamban R14del carriers: results of a large multicentre cohort study. Circ Cardiovasc Genet 2014;7:455–465.

14. Vafiadaki E, Glijnis PC, Doevendans PA, Kranias EG, Sanoudou D. Phospholamban R14del disease: The past, the present and the future. Frontiers in Cardiovascular Medicine 2023;10.

15. Nelson SED, Ha KN, Gopinath T, Exline MH, Mascioni A, Thomas DD, Veglia G. Effects of the Arg9Cys and Arg25Cys mutations on phospholamban’s conformational equilibrium in membrane bilayers. Biochimica et Biophysica Acta (BBA) - Biomembranes 2018;1860:1335–1341.

16. Schmitt JP, Ahmad F, Lorenz K, Hein L, Schulz S, Asahi M, Maclennan DH, Seidman CE, Seidman JG, Lohse MJ. Alterations of phospholamban function can exhibit cardiotoxic effects independent of excessive sarcoplasmic reticulum Ca2+-ATPase inhibition. Circulation 2009;119:436–444.

17. Kraev A. Insertional Mutagenesis Confounds the Mechanism of the Morbid Phenotype of a PLNR9C Transgenic Mouse Line. Journal of Cardiac Failure 2018;24:115–125.

18. Abrol N, de Tombe PP, Robia SL. Acute inotropic and lusitropic effects of cardiomyopathic R9C mutation of phospholamban. J Biol Chem 2015;290:7130–7140.

19. Ha KN, Masterson LR, Hou Z, Verardi R, Walsh N, Veglia G, Robia SL. Lethal Arg9Cys phospholamban mutation hinders Ca2+-ATPase regulation and phosphorylation by protein kinase A. Proc Natl Acad Sci U S A 2011;108:2735–2740.

20. Ceholski DK, Turnbull IC, Kong C-W, Koplev S, Mayourian J, Gorski PA, Stillitano F, Skodras AA, Nonnenmacher M, Cohen N, Björkegren JLM, Stroik DR, Cornea RL, Thomas DD, Li RA, Costa KD, Hajjar RJ. Functional and transcriptomic insights into pathogenesis of R9C phospholamban mutation using human induced pluripotent stem cell-derived cardiomyocytes. Journal of Molecular and Cellular Cardiology 2018;119:147–154.

21. Barndt RJ, Ma N, Tang Y, Haugh MP, Alamri LS, Chan SY, Wu H. Modeling of dilated cardiomyopathy by establishment of isogenic human iPSC lines carrying phospholamban C25T (R9C) mutation (UPITTi002-A-1) using CRISPR/Cas9 editing. Stem Cell Research 2021;56:102544.

22. Lian X, Hsiao C, Wilson G, Zhu K, Hazeltine LB, Azarin SM, Raval KK, Zhang J, Kamp TJ, Palecek SP. Robust cardiomyocyte differentiation from human pluripotent stem cells via temporal modulation of canonical Wnt signaling. Proceedings of the National Academy of Sciences 2012;109:E1848–E1857.

23. Ceholski DK, Trieber CA, Holmes CF, Young HS. Lethal, hereditary mutants of phospholamban elude phosphorylation by protein kinase A. J Biol Chem 2012;287:26596–26605.

24. Feyen DAM, McKeithan WL, Bruyneel AAN, Spiering S, Hörmann L, Ulmer B, Zhang H, Briganti F, Schweizer M, Hegyi B, Liao Z, Pölönen RP, Ginsburg KS, Lam CK, Serrano R, Wahlquist C, Kreymerman A, Vu M, Amatya PL, Behrens CS, Ranjbarvaziri S, Maas RGC, Greenhaw M, Bernstein D, Wu JC, Bers DM, Eschenhagen T, Metallo CM, Mercola M. Metabolic Maturation Media Improve Physiological Function of Human iPSC-Derived Cardiomyocytes. Cell Rep 2020;32:107925.

25. Barndt RJ, Liu Q, Tang Y, Haugh MP, Cui J, Chan SY, Wu H. Metabolic Maturation Exaggerates Abnormal Calcium Handling in a Lamp2 Knockout Human Pluripotent Stem Cell-Derived Cardiomyocyte Model of Danon Disease. Biomolecules 2023;13:69.

26. Nakagawa T, Yokoe S, Asahi M. Phospholamban degradation is induced by phosphorylation-mediated ubiquitination and inhibited by interaction with cardiac type Sarco(endo)plasmic reticulum Ca2+-ATPase. Biochemical and Biophysical Research Communications 2016;472:523–530.

27. Nickless A, Bailis JM, You Z. Control of gene expression through the nonsense-mediated RNA decay pathway. Cell & Bioscience 2017;7:26.

28. Tanida I, Ueno T, Kominami E. LC3 conjugation system in mammalian autophagy. The International Journal of Biochemistry & Cell Biology 2004;36:2503–2518.

29. Saito T, Sadoshima J. Molecular Mechanisms of Mitochondrial Autophagy/Mitophagy in the Heart. Circulation Research 2015;116:1477–1490.

30. Meng S, Cao J, He Q, Xiong L, Chang E, Radovick S, Wondisford FE, He L. Metformin activates AMP-activated protein kinase by promoting formation of the αβγ heterotrimeric complex. J Biol Chem 2015;290:3793–3802.

31. Sarkar S, Ravikumar B, Floto RA, Rubinsztein DC. Rapamycin and mTOR-independent autophagy inducers ameliorate toxicity of polyglutamine-expanded huntingtin and related proteinopathies. Cell Death & Differentiation 2009;16:46–56.

32. Vafiadaki E, Haghighi K, Arvanitis DA, Kranias EG, Sanoudou D. Aberrant PLN-R14del Protein Interactions Intensify SERCA2a Inhibition, Driving Impaired Ca2+ Handling and Arrhythmogenesis. International Journal of Molecular Sciences 2022;23:6947.

33. Raad N, Bittihn P, Cacheux M, Jeong D, Ilkan Z, Ceholski D, Kohlbrenner E, Zhang L, Cai C-L, Kranias EG, Hajjar RJ, Stillitano F, Akar FG. Arrhythmia Mechanism and Dynamics in a Humanized Mouse Model of Inherited Cardiomyopathy Caused by Phospholamban R14del Mutation. Circulation 2021;144:441–454.

34. Rijsingen IAWv, Zwaag PAvd, Groeneweg JA, Nannenberg EA, Jongbloed JDH, Zwinderman AH, Pinto YM, Deprez RHLd, Post JG, Tan HL, Boer RAd, Hauer RNW, Christiaans I, Berg MPvd, Tintelen JPv, Wilde AAM. Outcome in Phospholamban R14del Carriers. Circulation: Cardiovascular Genetics 2014;7:455–465.

35. Yang X, Rodriguez ML, Leonard A, Sun L, Fischer KA, Wang Y, Ritterhoff J, Zhao L, Kolwicz SC, Jr., Pabon L, Reinecke H, Sniadecki NJ, Tian R, Ruohola-Baker H, Xu H, Murry CE. Fatty Acids Enhance the Maturation of Cardiomyocytes Derived from Human Pluripotent Stem Cells. Stem Cell Reports 2019;13:657–668.

36. Lohse MJ, Engelhardt S, Eschenhagen T. What Is the Role of β-Adrenergic Signaling in Heart Failure? Circulation Research 2003;93:896–906.

37. Bristow MR. β-Adrenergic Receptor Blockade in Chronic Heart Failure. Circulation 2000;101:558–569.

38. Levine B, Kroemer G. Autophagy in the Pathogenesis of Disease. Cell 2008;132:27–42.

39. Nishida K, Kyoi S, Yamaguchi O, Sadoshima J, Otsu K. The role of autophagy in the heart. Cell Death & Differentiation 2009;16:31–38.

40. Zech ATL, Singh SR, Schlossarek S, Carrier L. Autophagy in cardiomyopathies. Biochimica et Biophysica Acta (BBA) - Molecular Cell Research 2020;1867:118432.

41. Taneike M, Yamaguchi O, Nakai A, Hikoso S, Takeda T, Mizote I, Oka T, Tamai T, Oyabu J, Murakawa T, Nishida K, Shimizu T, Hori M, Komuro I, Takuji Shirasawa TS, Mizushima N, Otsu K. Inhibition of autophagy in the heart induces age-related cardiomyopathy. Autophagy 2010;6:600–606.

42. Bhuiyan MS, Pattison JS, Osinska H, James J, Gulick J, McLendon PM, Hill JA, Sadoshima J, Robbins J. Enhanced autophagy ameliorates cardiac proteinopathy. The Journal of Clinical Investigation 2013;123:5284–5297.

43. Gupta MK, McLendon PM, Gulick J, James J, Khalili K, Robbins J. UBC9-Mediated Sumoylation Favorably Impacts Cardiac Function in Compromised Hearts. Circulation Research 2016;118:1894–1905.

44. Marin TM, Keith K, Davies B, Conner DA, Guha P, Kalaitzidis D, Wu X, Lauriol J, Wang B, Bauer M, Bronson R, Franchini KG, Neel BG, Kontaridis MI. Rapamycin reverses hypertrophic cardiomyopathy in a mouse model of LEOPARD syndrome–associated PTPN11 mutation. The Journal of Clinical Investigation 2011;121:1026–1043.

45. Xie Z, Lau K, Eby B, Lozano P, He C, Pennington B, Li H, Rathi S, Dong Y, Tian R, Kem D, Zou M-H. Improvement of Cardiac Functions by Chronic Metformin Treatment Is Associated With Enhanced Cardiac Autophagy in Diabetic OVE26 Mice. Diabetes 2011;60:1770–1778.

46. Dai D-F, Karunadharma PP, Chiao YA, Basisty N, Crispin D, Hsieh EJ, Chen T, Gu H, Djukovic D, Raftery D, Beyer RP, MacCoss MJ, Rabinovitch PS. Altered proteome turnover and remodeling by short-term caloric restriction or rapamycin rejuvenate the aging heart. Aging Cell 2014;13:529–539.

47. Eisenberg T, Abdellatif M, Schroeder S, Primessnig U, Stekovic S, Pendl T, Harger A, Schipke J, Zimmermann A, Schmidt A, Tong M, Ruckenstuhl C, Dammbrueck C, Gross AS, Herbst V, Magnes C, Trausinger G, Narath S, Meinitzer A, Hu Z, Kirsch A, Eller K, Carmona-Gutierrez D, Büttner S, Pietrocola F, Knittelfelder O, Schrepfer E, Rockenfeller P, Simonini C, Rahn A, Horsch M, Moreth K, Beckers J, Fuchs H, Gailus-Durner V, Neff F, Janik D, Rathkolb B, Rozman J, de Angelis MH, Moustafa T, Haemmerle G, Mayr M, Willeit P, von Frieling-Salewsky M, Pieske B, Scorrano L, Pieber T, Pechlaner R, Willeit J, Sigrist SJ, Linke WA, Mühlfeld C, Sadoshima J, Dengjel J, Kiechl S, Kroemer G, Sedej S, Madeo F. Cardioprotection and lifespan extension by the natural polyamine spermidine. Nature Medicine 2016;22:1428–1438.

48. Singh SR, Zech ATL, Geertz B, Reischmann-Düsener S, Osinska H, Prondzynski M, Krämer E, Meng Q, Redwood C, Velden Jvd, Robbins J, Schlossarek S, Carrier L. Activation of Autophagy Ameliorates Cardiomyopathy in *Mybpc3*-Targeted Knockin Mice. Circulation: Heart Failure 2017;10:e004140.

49. Feyen DAM, Perea-Gil I, Maas RGC, Harakalova M, Gavidia AA, Arthur Ataam J, Wu TH, Vink A, Pei J, Vadgama N, Suurmeijer AJ, Te Rijdt WP, Vu M, Amatya PL, Prado M, Zhang Y, Dunkenberger L, Sluijter JPG, Sallam K, Asselbergs FW, Mercola M, Karakikes I. Unfolded Protein Response as a Compensatory Mechanism and Potential Therapeutic Target in PLN R14del Cardiomyopathy. Circulation 2021;144:382–392.

50. Kumar M, Haghighi K, Koch S, Rubinstein J, Stillitano F, Hajjar RJ, Kranias EG, Sadayappan S. Myofilament Alterations Associated with Human R14del-Phospholamban Cardiomyopathy. Int J Mol Sci 2023;24.

51. Teng ACT, Miyake T, Yokoe S, Zhang L, Rezende LM, Sharma P, MacLennan DH, Liu PP, Gramolini AO. Metformin increases degradation of phospholamban via autophagy in cardiomyocytes. Proceedings of the National Academy of Sciences 2015;112:7165–7170.

52. Li J, Kim SG, Blenis J. Rapamycin: one drug, many effects. Cell Metab 2014;19:373–379.

53. Miller RA, Birnbaum MJ. An energetic tale of AMPK-independent effects of metformin. The Journal of Clinical Investigation 2010;120:2267–2270.

